# Transcript quantification using context-aware MAPQ modeling

**DOI:** 10.64898/2026.10.26.761703

**Authors:** Sowmya Parthiban, Stephanie C. Hicks

## Abstract

Long-read RNA-sequencing enables transcript-level quantification, but filtering alignments using fixed mapping quality (MAPQ) thresholds can discard reads reflecting intrinsic transcriptome ambiguity and distort abundance estimates. Here we present CALMeR (Context-Aware Long-read MAPQ-based Rescuer), a regression-based approach that fits transcript and read features. By modeling the expected mapping quality of each alignment, the method distinguishes low scores consistent with transcript context from anomalously low scores indicative of artifacts. This approach retains reads from multi-isoform genes and paralog families that would otherwise be excluded, while removing alignments with unexpectedly low confidence. Retained reads can be counted per transcript or supplied to downstream quantification tools. In spike-in benchmarks, context-aware filtering recovers more transcripts and yields abundance estimates that better match known concentrations than a MAPQ ≥ 1 filter and expectation-maximization-based quantification methods. These results support incorporating transcript context into alignment filtering for long-read RNA sequencing. Open-source software is available at https://github.com/sparthib/CALMeR.

## 1 Introduction

Multi-mapping reads pose a fundamental challenge for RNA quantification, as they cannot be unambiguously assigned to a single transcript or genomic location. The MAPping Quality (MAPQ) score — defined in the SAM specification as −10 × log_10_(Pr{mapping position is wrong}) rounded to the nearest integer [1] — provides a Phred-scaled measure of alignment confidence or the probability that the mapping position is incorrect. For example, in `minimap2` [2], an aligner developed specifically for long-read sequencing, MAPQ score ranges from 0 to 60. A MAPQ of 30, 20, and 10 corresponds to 99.9%, 99%, and 90% confidence (or 1 in 1000, 1 in 100, 1 in 10 chance of incorrect mapping), respectively (**Figure S1**). In contrast, the interpretation of a MAPQ score of 0 is that it is equally likely for the read to map to two or more locations. However, there are several challenges with using the MAPQ score to select reads for quantification in practice. First, MAPQ is computed by each aligner using its own empirical heuristics, so the same numerical score carries different meanings across tools and cannot be compared between them [3]. Second, its interpretation depends on the choice of reference. When reads are aligned to a genome, all isoforms of a gene project onto a single genomic locus, and a low MAPQ implies ambiguity across loci — paralogs, pseudogenes, or repeats. When reads are aligned to a transcriptome, every annotated isoform is a distinct reference sequence, and a low MAPQ could additionally imply ambiguity across isoforms of the same gene as well as across loci [4, 5]. Third, there is a lack of principled basis for the static threshold used to filter reads in downstream analysis, and the fraction of reads discarded by any fixed threshold varies unpredictably across genes depending on local sequence complexity and isoform structure [4, 6]. Finally, removing ambiguous reads can carry huge quantification cost: in short-read isoform quantification, methods that probabilistically model multi-mapping consistently outperform count-based approaches that discard such reads [7].

Existing long-read quantification tools take markedly different stances on whether, and how, to use MAPQ as a filter for alignment quality. `FLAIR` [8], a long-read isoform discovery and quantification tool, explicitly leverages MAPQ as a metric for filtering alignments for quantification. After realigning reads to its first-pass isoform set, the authors of FLAIR benchmarked three quantification strategies on SIRV controls and simulated nanopore reads — primary alignments only, alignments with MAPQ ≥ 1, and `salmon` in alignment-based mode — and adopted MAPQ ≥ 1 as its default for quantification because it correlated best with expected counts. The authors note, however, that this approach discards a substantial fraction of data: 25.5% of reads (5,469 of 21,456) in the SIRV benchmark had MAPQ = 0 and were excluded from quantification, motivating their inclusion of `salmon` [9] as an optional alternative. On the other hand, methods such as `oarfish` [10], `salmon` [9], and `Bambu` [11] apply no MAPQ filter at all, instead using an Expectation-Maximization approach that probabilistically distributes each read’s weight across all of its candidate transcripts.

This split in practice — MAPQ ≥ 1 filtering on one side, no MAPQ filtering on the other — points to a missing third option for quantification. Rather than treating MAPQ as a blanket filter or omitting it altogether, which forfeits a useful signal of alignment quality, we propose selecting reads for quantification by dynamically filtering on MAPQ. Under this approach, a read’s observed MAPQ is compared against what its transcript and read context would predict: reads that deviate are flagged and removed, while reads whose MAPQ is consistent with their context are kept and counted for a given transcript. Crucially, this preserves reads with context-consistent low MAPQ — those arising from multi-isoform genes, paralog families, and other transcript features — rather than discarding them as a fixed threshold would.

Here we show that a read’s MAPQ depends systematically on transcript and read properties: longer transcripts face systematic under-sampling due to pore blockage or inefficient reverse transcription, and their MAPQ scores fluctuate with GC content and isoform complexity. These dependencies are precisely what make a single fixed threshold problematic for quantification, and what make a model-based read-selection approach worth investigating further. Such an approach requires a model of how MAPQ depends on transcript and read properties — and the long-read field is unusually well-positioned to build one. Unlike short-read RNA-seq, where multiple aligners (`BWA-MEM` [12], `Bowtie2` [13], and others) implement MAPQ with different conventions, long-read isoform analysis has converged on `minimap2` as a widely-used aligner [2, 14–16]. This commonality makes it tractable to characterize how alignment MAPQ associates with transcript-level features across long-read studies.

Here, we present `CALMeR` (Context-Aware Long-read MAPQ-based Rescuer), a method that selects reads for transcript quantification by learning, per sample, how MAPQ depends on transcript- and read-level features. `CALMeR` identifies reads whose MAPQ is anomalously low relative to that learned context, and provides a filtered BAM in which biology-driven ambiguity is preserved while context-anomalous noise is removed - so that the retained reads can be counted per transcript for quantification. We show that quantifying from the reads `CALMeR` retains recovers reads from multi-isoform genes and paralog families that fixed MAPQ thresholds discard, and yields transcript abundances that agree more closely with spike-in ground truth than either MAPQ ≥ 1 filtering or no-filter EM quantification. `CALMeR` is implemented as an R package with diagnostic visualizations of MAPQ–feature relationships, allowing users to inspect the conditional model before implementing it.

## 2 Results

### 2.1 Datasets

We identified five publicly available datasets from recent benchmarking studies, spanning direct RNA, direct cDNA, PCR cDNA, R2C2, and CapTrap-Seq library preparations across 8 kits (**Table S1**). The five datasets stem from (i) Dong et al. [17], (ii) Long-read RNA-Seq Genome Annotation Assessment Project (LRGASP) [14], (iii) LongBench [16], (iv) Singapore Nanopore Expression (SG-NEx) Project [15], and (v) Carbonell-Sala et al. [18] who evaluate CapTrap-Seq against nanopore direct RNA and other non-nanopore protocols. These datasets contain synthetic spike-ins, including Lexogen SIRVs [19], sequins [20], or a combination of both, with composition and concentration varying by sample (**Table S2**). We uniformly processed all samples (**Methods**) to benchmark performance across legacy and current kits. More specifically, we focused on PCR cDNA kits because they have been shown to be most prone to coverage breaks reported in previous benchmarking studies [15, 16], which degrade mapping confidence. We used these data to examine how MAPQ scores from long-read RNA-seq reads aligned with `minimap2` vary with transcript-level features and evaluating the impact of read and alignment filtering strategies on quantification.

### 2.2 MAPQ scores vary as a function of transcript-level features

To examine how MAPQ varies with transcript-level features, we computed the average MAPQ across aligned reads considering four properties of each transcript: (i) the number of isoforms in its parent gene, (ii) GC content, (iii) transcript length, and (iv) the fraction of the transcript covered by the alignment (**Figure 1**). Specifically, we analyzed PCS109 samples from Dong et al. [17] and SG-NEx [15], PCS110 from LRGASP [14], and PCS114 from LongBench [16] (oldest to newest PCR cDNA kits), for detecting changes in MAPQ trends between legacy and current kits.

**Figure 1:**
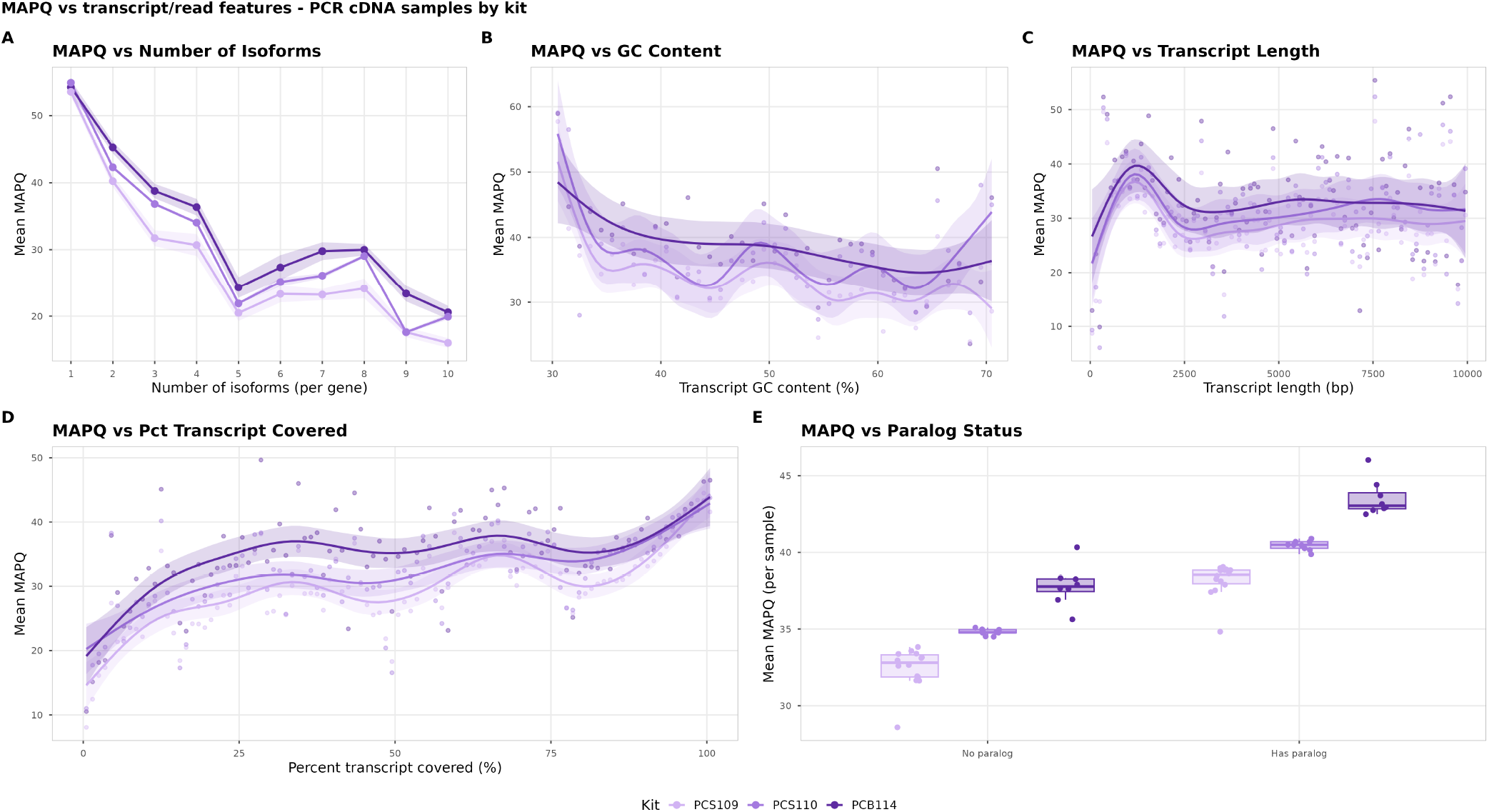
MAPQ scores vary with transcript-level features. We focused on samples generated with three nanopore PCR cDNA primer (or ligation) kits: PCS109 [15, 17], PCS110 [14], and PCS114 [16] (oldest to newest, respectively). In panels A–D, each point represents the per-sample mean MAPQ within a bin of the indicated transcript feature; lines and ribbons summarise across samples within each kit. (**A**) Mean MAPQ versus the number of isoforms annotated for the parent gene (1–10), shown as a kit-specific line ± 95% CI ribbon over per-sample bin means. (**B**) Mean MAPQ versus transcript GC content (1% bins, 30–70%). (**C**) Mean MAPQ versus reference transcript length (100 base pair (bp) bins, ≤ 10,000 bp). (**D**) Mean MAPQ versus the percent of the reference transcript length covered by the read (1% bins). In panels B, C, and D, per-sample bin means are overlaid with a kit-specific Generalized Additive Models (GAM) smoother and its 95% confidence band. (**E**) Distribution of per-sample mean MAPQ on transcripts with versus without a paralog, shown as kit-coloured boxplots with overlaid sample points; paralogue status is the binary indicator produced by the CALMeR self-alignment pipeline (**Methods**).

Across all kits, we identify several key findings. First, average MAPQ across samples decreases in a non-linear fashion with a steeper downward slope from *n*_iso_ = 1 to *n*_iso_ = 5 and declines slower with more multi-isoformic genes (**Figure 1A**). Second, the average MAPQ mildly decreases with increasing transcript GC content (**Figure 1B**). Third, average MAPQ across samples is extremely low for reads mapping to transcripts < 500 base pairs (bp), but is the highest at 500bp, and decreases as transcript length increases to 2500bp and plateaus (**Figure 1C**). Lastly, average MAPQ increases with the percent of the reference transcript length covered by the alignment, since reads that span more of their target transcript provide more unique sequence context and reduce competing secondary alignments) (**Figure 1D**). Baseline MAPQ values tend to go higher with newer kits, with PCS114 being the latest, and PCS109 the oldest legacy nanopore kit. Surprisingly, transcripts with no detectable paralogs in other genes have lower MAPQ, on average, than transcripts that do have paralogs (**Figure 1E**).

However, stratifying by number of isoforms reveals that within every isoform bin from *n*_iso_ = 1 to *n*_iso_ = 10+, the negative transcript length–MAPQ relationship previously described (**Figure 1A**) flattens within each bin, but the overall mean within each *n*_iso_ stratification shifts downward (**Supplementary Figure S3A**). This suggests that multi-isoform loci decreases MAPQ independently of transcript length. In contrast, the relationship between MAPQ and percent covered is likewise modified by reference transcript length, dampening for longer transcripts above 5000 bp (**Supplementary Figure S3B**). Finally, we further investigated the relationship between MAPQ and paralog status (**Figure 1E**) by now stratifying by *n*_iso_. We found that within nearly every isoform stratum, reads on transcripts with a cross-gene paralog carry lower MAPQ than reads on transcripts without one (**Supplementary Figure S3C**) — reversing the marginal association shown above, where the raw split is confounded by *n*_iso_. Together, these observations expose the central limitation of static MAPQ thresholds: a uniform cutoff disproportionately discards reads from multi-isoform genes, GC-extreme transcripts, transcripts with cross-gene paralogs, and — within isoform strata — longer transcripts. The strong role of these interactions motivates a modeling framework that conditions on transcript-level context rather than applying a single MAPQ cutoff uniformly for all genes.

### 2.3 Static MAPQ filtering disproportionately under represents reads from multi-isoform genes

The analysis above characterizes the structural dependencies of MAPQ in the context of transcript-level features, but it does not directly demonstrate that these dependencies translate into measurable quantification bias - there is no ground-truth known abundance against which to compare observed counts. To obtain that direct measurement, we turn to spike-ins, where the input concentrations of each transcript are known by construction and any deviation between observed and expected proportions can be attributed to upstream technical bias.

Here, we focus on a set of PCR cDNA samples (*n* = 8) with Sequins Mix A v1 [21] used in the Dong et al. [17] and You et al. [16] datasets (**Figure 2**). We picked these samples because they span the broadest dynamic range of the relevant transcript-level features among the available spike-in panels: isoform count per gene 1–4, transcript length 283 bp to 6943 bp, and GC content 32.5–68.9%. For a given static MAPQ threshold *α* (with *α* = 0 corresponding to no filtering), we calculated the per-transcript log_2_-fold change between the observed and expected within-panel proportion of each spike-in transcript *t* as

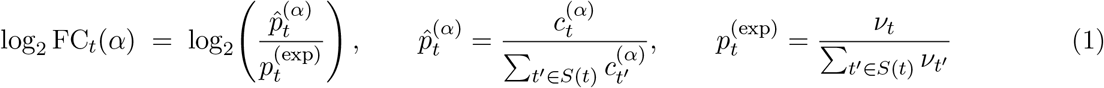

where 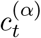 is the number of primary alignments aligned to transcript *t* after retaining only reads with MAPQ ≥ *α, ν*_*t*_ is the manufacturer-supplied nominal concentration for *t*, and *S*(*t*) is the spike-in panel containing *t* (here, Sequins Mix A v1). Normalising within the panel makes the units of *c* and *ν* (read counts vs. nominal mix concentration) cancel.

**Figure 2:**
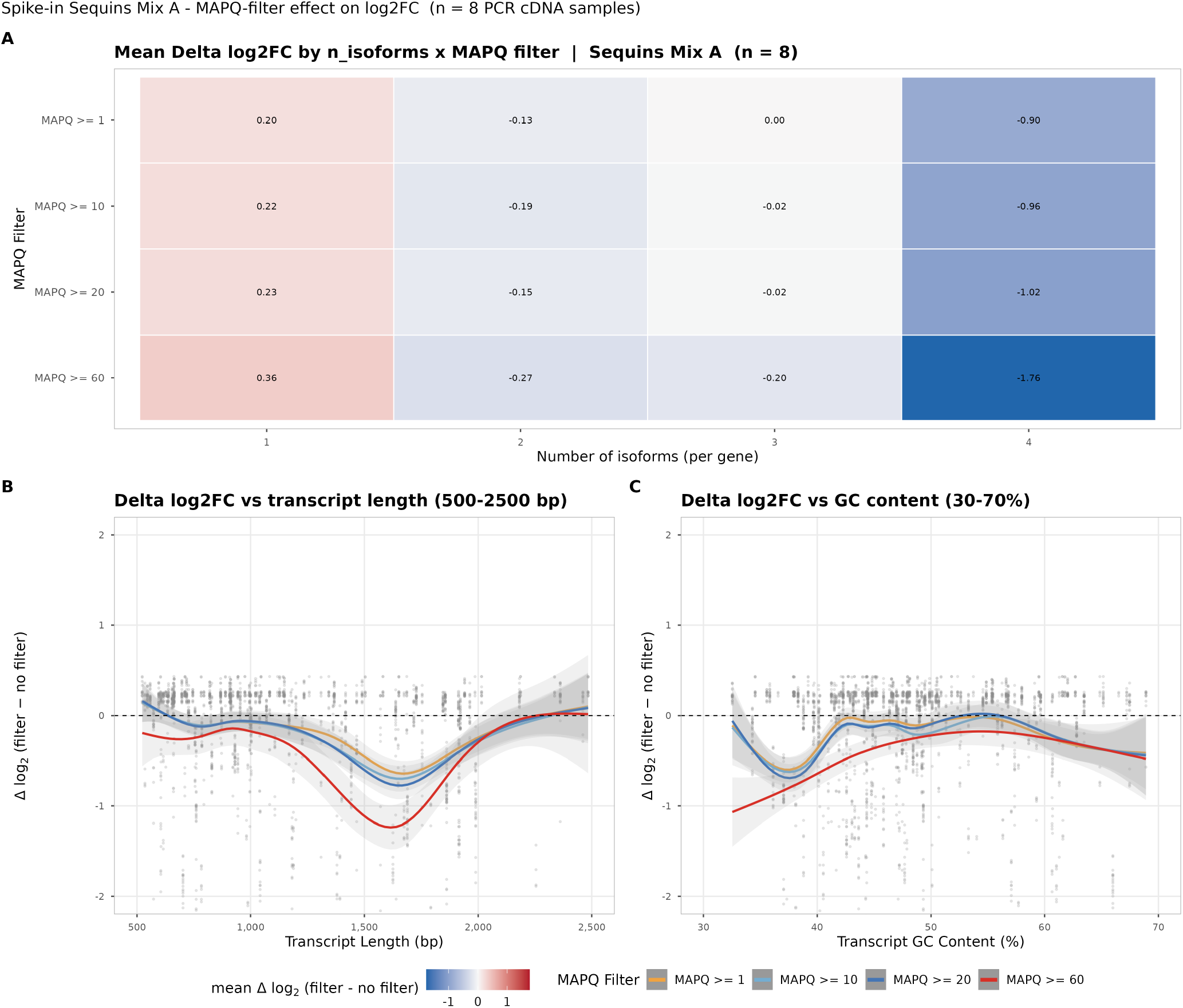
Hard MAPQ filtering disproportionately underrepresents reads from multi isoform genes. (**A**) Mean Δ log_2_ FC per (number of annotated isoforms per gene) × (MAPQ filter), across *n* = 8 PCR cDNA samples carrying Sequins Mix A (PCS109 from Dong et al. and PCS114 from Longbench). Negative cells (blue) indicate under representation relative to the no-MAPQ-filter baseline. (**B**) Δ log_2_ FC as a function of transcript length, restricted to 500–2500 bp where the bulk of spike-in reads concentrate; GAM smooths ± 95% CI, colored by MAPQ filter. (**C**) Same, versus transcript GC content (30–70%). Δ log_2_ FC = log_2_ FC(filter) − log_2_ FC(no MAPQ filter).

We then quantified the difference between log_2_ FC_*t*_(*α*) retaining only reads with MAPQ ≥ *α* compared to a baseline log2 FC_*t*_(*α* = 0) (no MAPQ filtering):

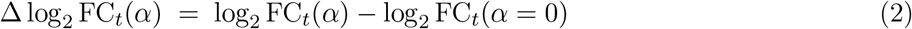

In a setting where there is no quantification bias, we would expect Δ log_2_ FC = 0. However, we found that single-isoform genes (*n*_iso_ = 1) are mildly overrepresented under the most liberal filter (MAPQ ≥ 1) and increase in over representation with stricter cutoffs (**Figure 2A**). On the other hand, *n*_iso_ = 4 is consistently underrepresented, and the magnitude of under representation grows with stringency. In contrast, transcripts in the *n*_iso_ = 2 and *n*_iso_ = 3 categories do not show meaningful changes across cutoffs. This supports our earlier findings based on MAPQ scores varying as a function of transcript-level features (**Figure 1**), namely that static MAPQ thresholding biases towards keeping reads mapping to single-isoform genes and biases against multi-isoform genes. In addition to investigating how the number of isoforms impacts quantification bias, we also explore how transcript length and GC content impact quantification bias. We found that Δ log_2_ FC becomes increasingly negative with transcript length up to ∼1700 bp and then reverses (**Figure 2B**). GC content shows no strong overall trend across most of the 30–70% range (**Figure 2C**).

However, these observations should be cautiously interpreted. When comparing Sequins Mix A feature distributions against GENCODE v46, it is worth noting that the sequin transcripts span a substantially narrower length range than endogenous transcripts (median 1,076 bp vs. 2,297 bp for protein-coding transcripts; only 0.6% of sequins exceed 5 kb, against 14.5% in the protein-coding subset). Although sequins are deliberately multi-isoform by design — 88.2% of sequin genes carry ≥ 2 annotated isoforms, exceeding the 66.6% protein-coding rate—the design caps at four isoforms per gene and therefore severely under-samples the highly multi-isoform loci that drive the strongest filtering biases at scale: only 3.9% of Sequins Mix A genes carry four or more annotated isoforms, compared with 10.9% across the full GTF and 27.9% within the protein-coding subset alone (**Figure S2A**). Even though sequins extend beyond the SIRV panel, their dynamic range still falls well short of the endogenous distribution. Sequins also tend to be shorter than bioligical transcripts, especially those of protein coding genes (**Figure S2B**). The GC distributions overlap broadly between sequins and the GENCODE protein-coding subset (median 47.3% vs. 49.2%; comparable IQR) (**Figure S2C**). However, prior work has noted that sequins lack the structural complexity of native transcripts [14, 15, 22]. We therefore treat sequins as a complementary measurement to the transcriptome-wide MAPQ trends.

Together, these observations motivate a model-based filtering approach in which MAPQ is interpreted conditional on transcript and read-level context. The four transcript-level features — *n*_isoforms_, GC content, transcript length, and a binary indicator of whether the isoform has a paralog (has_paralog; defined in **Methods**, “Paralog Detection”) — are joined by one read-level feature: the percent of the transcript that the alignment covers. While base calling quality is controlled upstream at *Q >* 10, percent transcript covered captures a complementary axis of read informativeness—a long, high-quality read spanning a greater proportion of its target transcript provides more unique sequence context and further reduces mapping ambiguity (**Figure 1D**).

### 2.4 Beta sampling distribution for modeling MAPQ scores

As noted in the introduction, the MAPQ score for each read ranges between 0 and 60 (**Figure S1**). To explore the distribution of reads, we randomly sampled 50,000 primary aligned reads (secondary and supplementary read alignments excluded) from three samples across three sequencing protocols, including PCR cDNA, direct RNA, and direct cDNA. This includes the PCS114 sample from You et al. [16] (H1975 cell line; accession GSM9135480), which will be used throughout for model development and per-read scoring in the following sections.

For a given read *i*, we consider the original MAPQ score for read *i* as *z*_*i*_ ∈ (0, 60). We found that there is a substantial fraction of primary aligned reads with *z*_*i*_ = 0 (ambiguous read alignment quality) and *z*_*i*_ = 60 (unique read alignment quality) (**Figure S4**). This motivated us to define a rescaled version 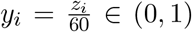 so that we could model the MAPQ scores with a beta regression model [23]. However, the rescaled MAPQ response *y*_*i*_ ∈ [0, 1] presents a modeling challenge that standard beta regression alone cannot accommodate. Beta regression is the canonical choice for continuous proportions because the beta density is flexible across [0, 1] — a two-parameter family that can take left-skewed, right-skewed, U-shaped, inverted-U, or uniform shapes depending on its parameters. The crucial limitation, however, is that the beta distribution is purely continuous: it assigns *zero* probability to any individual point in the interval. In our setting a non-trivial fraction of primary alignments sit at the exact boundaries — *y*_*i*_ = 0 (MAPQ = 0, fully ambiguous mappings) and *y*_*i*_ = 1 (MAPQ = 60, maximum aligner-assigned uniqueness) — so a beta-only specification cannot describe the empirical response at all without ad-hoc nudging away from {0, 1}. We therefore adopt the zero-and-one inflated beta (ZOIB) regression model of Ospina and Ferrari [24], in which the response is modeled as a three-component mixture: a beta density on the open interior (0, 1) together with two Bernoulli-type point masses at *y*_*i*_ = 0 and *y*_*i*_ = 1. Each of the three component parameters (the beta mean and precision, plus the two boundary-mass probabilities) is allowed to depend on the predictor vector through its own sub-model, yielding a single jointly estimated regression for a mixed continuous–discrete response.

### 2.5 CALMeR identifies low quality mapping reads using outlier detection

Our proposed framework CALMeR models the rescaled MAPQ scores (*y*_*i*_) as a function of the five transcript- and read-level features (**x**_*i*_) using a ZOIB regression model [24] and estimated the `gamlss` framework [25] with *θ* = (***β, γ, δ***) denoting a coefficient vector for the interior, zero-inflation, and one-inflation sub-models, respectively (**Methods**). In contrast to using static MAPQ thresholds to remove low-quality mapping reads, we identify reads with low mapping quality in terms of an outlier detection framework. Specifically for each read alignment, we propose to use a randomized quantile residual (RQR) [26]:

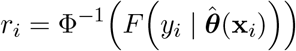

for interior values *y*_*i*_ ∈ (0, 1) that are evaluated using the fitted cumulative distribution function (CDF) 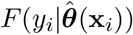 and finding the corresponding quantile of the standard normal distribution Φ^−1^(·). For discrete outcomes, a random value is drawn uniformly between the lower and upper limits of the CDF jump interval before applying the inverse normal transformation to avoid “jumps” or steps in the CDF. Under correct model specification, *r*_*i*_ is approximately standard normal, so deviations from *N* (0, 1) jointly summarize model adequacy and per-read anomaly. Alignments whose covariates fall outside the training support are returned as unscored rather than extrapolated.

Next, we describe the transcript- and read-level features (**x**_*i*_) annotated for each read’s primary alignment. Five covariates were considered: four transcript-level features (constant across all reads aligning to the same transcript) and one read-level feature (which depends on the individual alignment). The four transcript-level features are: (i) **transcript length** (tx_length), the total transcript length in nucleotides, computed as the sum of exon widths from the GTF annotation; (ii) **GC content** (gc_content), the proportion of G and C bases in the transcript sequence, obtained from the transcriptome FASTA and expressed as a percent; (iii) **number of isoforms** (n_isoforms), the count of annotated isoforms for the parent gene, obtained by counting distinct transcript identifiers per gene in the GTF; and (iv) **paralog presence** (has_paralog), a binary indicator for whether the target transcript has at least one cross-gene paralog passing the alignment-stage filters described in Methods. The read-level feature is (v) **percent covered** (pct_covered), the percent of the transcript covered (or the proportion of the transcript length spanned by the read alignment, computed as alignment_length*/*tx_length × 100 and capped at 100%). Note, the predictor distributions of the five covariates were modeled as either smoothed (using penalized B-splines on a standardized continuous variable), piecewise linear (an integer count with a kink at a fixed knot), or plain linear (a binary indicator) (**Figure S5**).

Using 50,000 reads randomly drawn from the PCB114/H1975 sample (GSM9135480), we defined the covariate structure for the ZOIB regression model of MAPQ in the CALMeR diagnostic framework. Term selection was carried out by backward elimination under the generalized Akaike information criterion (GAIC) with a BIC penalty applied across the distribution parameters using the two strategies implemented in the `gamlss` package [25] along with `stepGAICAll.A` and `stepGAICAll.B` [27]. Strategy A selects a possibly different subset of terms for each parameter (*µ, σ, ν, τ* ), whereas Strategy B selects a single set of terms shared by all parameters; both drop, at each step, the term whose removal most improves GAIC, subject to marginality. *σ* was held constant (*σ* ∼ 1), while *µ, ν*, and *τ* were modeled with the full covariate pool. Starting from the full model—penalized B-spline smooths for the three continuous covariates (transcript length, GC content, percent covered), a two-piece linear basis for isoform count, the binary `has_paralog` indicator, and the three transcript-feature interactions—backward elimination removed no terms under either strategy: both converged on the identical full model (global deviance 72,158.8 on 81.1 effective degrees of freedom; AIC 72,320.9; BIC 73,036.1; Nagelkerke’s *R*^2^ = 0.461), confirming that every covariate, in every parameter equation, lowers GAIC.

Using the same 50,000 reads from PCS114 / H1975 sample (GSM9135480), we found the full model with all 5 predictors and 3 interaction terms fit the data well as shown in the RQR quantile-quantile (QQ) plot (**Figure 3**). Specifically, the empirical quantiles track the theoretical *N* (0, 1) reference line closely across essentially the full range, indicating that the fitted ZOIB distribution captures both the bulk and the tails of the rescaled-MAPQ response on the training data.

**Figure 3:**
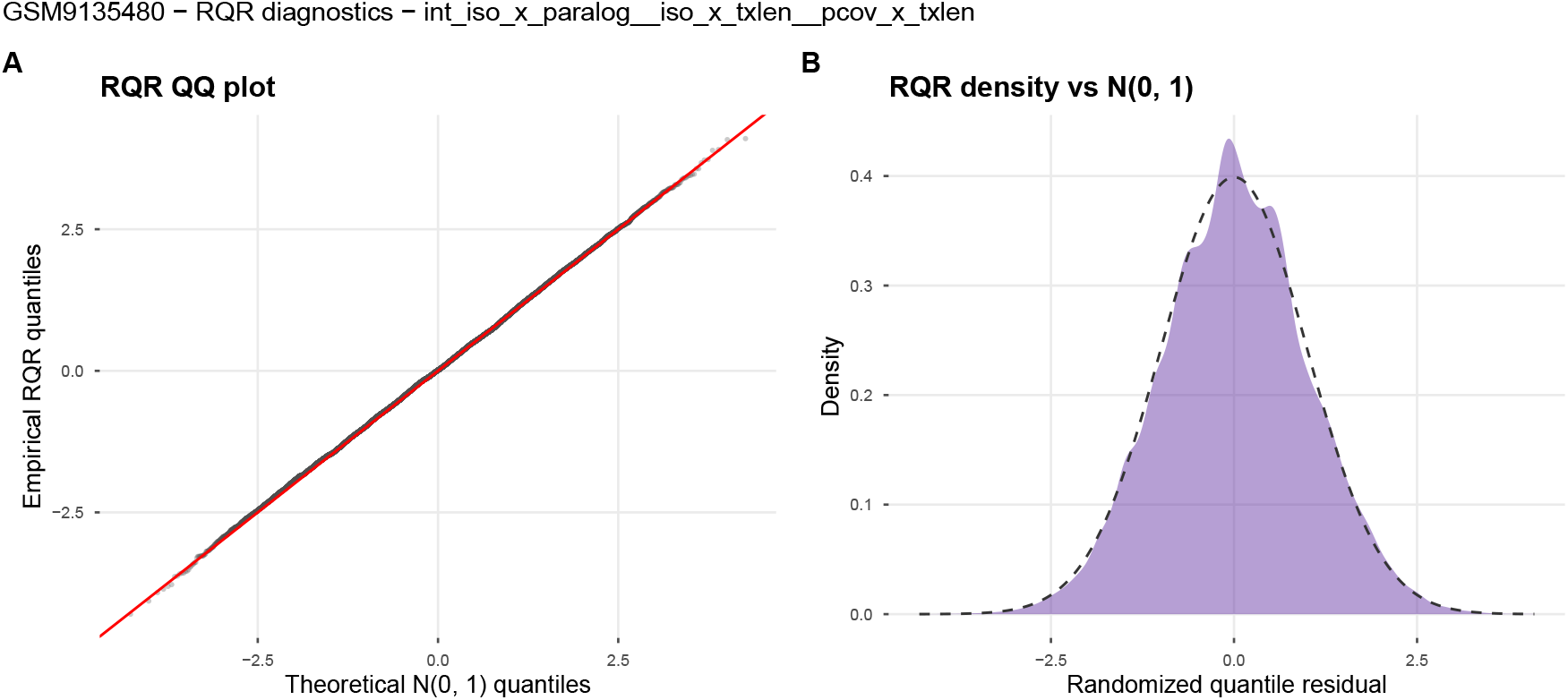
MAPQ scores modeled with a zero-and-one inflated beta (ZOIB) regression model. Randomized quantile residual (RQR) calculated using the full fitted model’s ZOIB cumulative distribution function on sample GSM9135480. (**A**) Diagnostic quantile-quantile (QQ) plot of the empirical RQR distribution against the theoretical *N* (0, 1) quantiles; the red reference line indicates the identity. (**B**) Density of the empirical RQR distribution (filled) overlaid on the N (0, 1) density (dashed). Both panels indicate that the fitted model captures the response distribution well across essentially the full quantile range.

Next, we evaluated how the model performed in terms of mapping each predictor onto the expected MAPQ. Holding the other predictors at their training-median values, the expected MAPQ appears to piecewise decrease and plateau as a function of `n_isoforms` (**Figure 4A**). The `gc_content` curve is approximately flat across the central range (GC ≲ 35% and GC ≳ 65%) and expected MAPQ peaks around `tx_length` ∼1,000 bp and falls sharply only past ∼10 kb (**Figure 4B,C**). The `pct_covered` panel increases non-linearly, and `has_paralog` shifts expected MAPQ downward by approximately 5 points when set to TRUE, in line with the higher mapping ambiguity expected for transcripts with a cross-gene paralog (**Figure 4D,E**). Together, these effects illustrate MAPQ score influenced by various transcript and gene-level.

**Figure 4:**
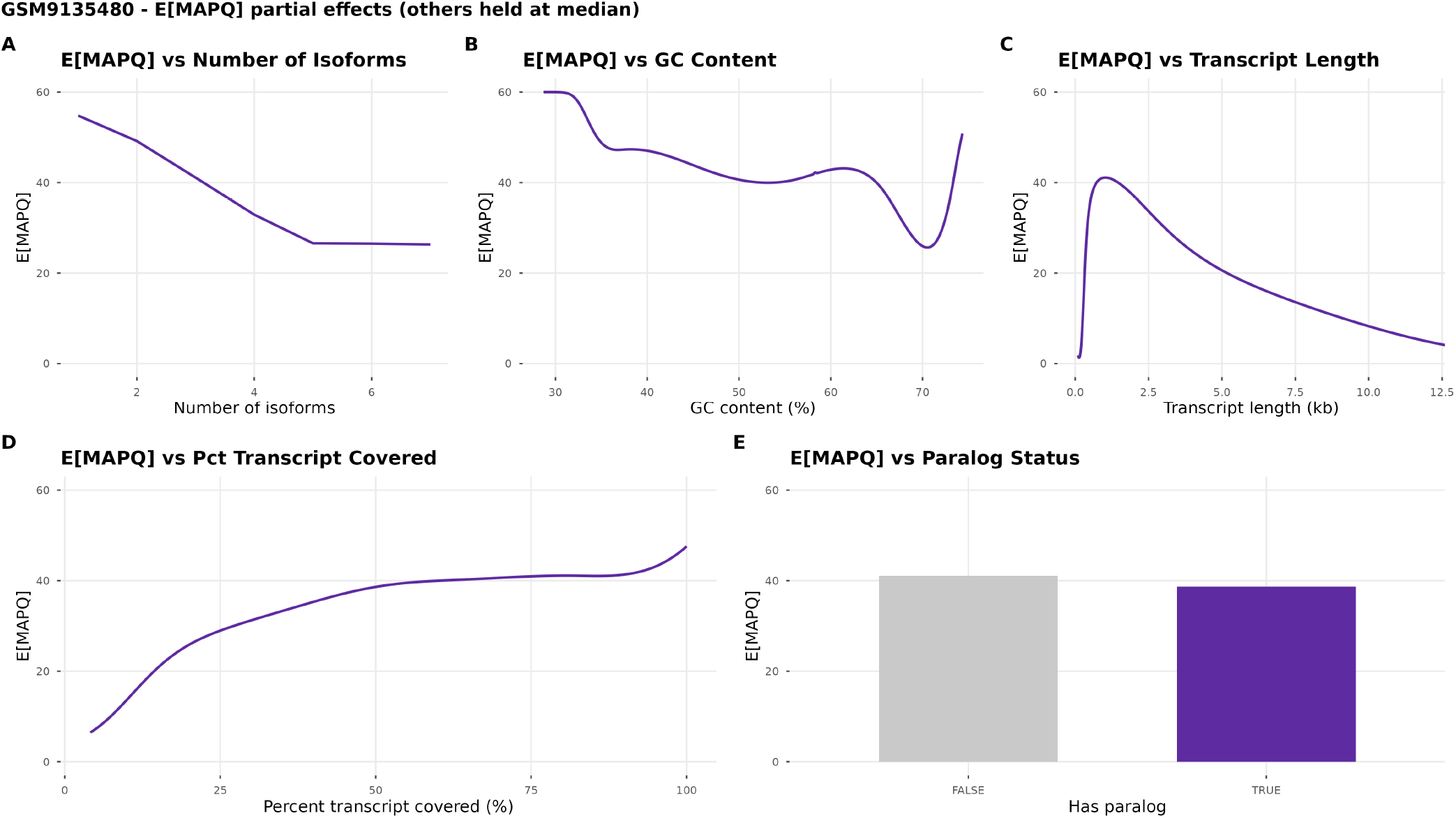
Partial effects of each predictor on expected MAPQ. Each panel varies one predictor across its training range while holding the other predictors at their median value (or `FALSE` for the binary `has_paralog` indicator), and plots E[MAPQ] = 60 · (*τ* + *µ*)*/*(1 + *ν* + *τ* ) under the resulting prediction. (**A**) `n_isoforms` is entered as a two-piece linear basis with a knot at 5. (**B-D**) Smoothed predictors `gc_content`, `tx_length` (log-bp *x*-axis), and `pct_covered` are entered through penalized B-splines and appear as continuous curves. (**E**) `has_paralog` is binary and shown as a column comparison.

To identify low quality mapping reads, we applied a filter of two median absolute deviations (MAD), which identified a small, consistent fraction of scorable primary alignments—1.31%, 1.56%, and 1.16% in H1975, HCC827, and H211 samples, respectively (**Figure 5**). Flagged and retained reads separated most sharply in mapping quality: flagged reads had a median MAPQ of ∼1 versus 46–52 for retained reads (**Figure 5A**). Also, flagged reads sat on modestly shorter transcripts (Cohen’s *d* ≈ 0.5), but they had essentially identical GC content (median ≈ 50% in both flagged and unflagged; *d* ≈ 0.04 (**Figure 5B,C**).

**Figure 5:**
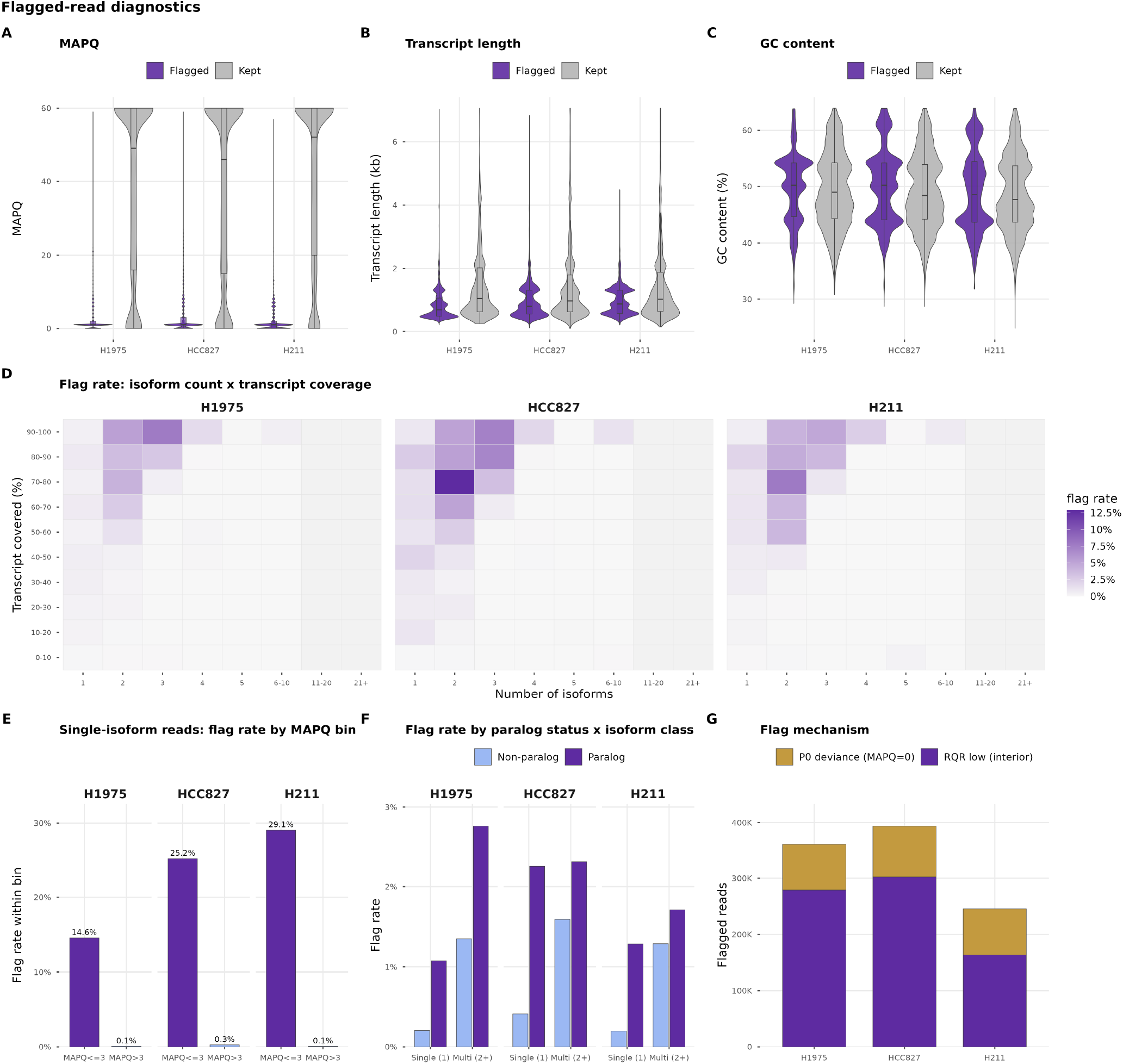
Reads flagged by CALMeR are low-MAPQ, high-coverage alignments enriched at paralogous and multi-isoform transcripts. Diagnostics contrasting reads flagged by the 2-MAD rule against retained reads, among scorable primary alignments, across three PCS114 spike-in samples (H1975, HCC827, H211). (**A–C**) Split violins (flagged, purple; retained, grey) for MAPQ, transcript length, and GC content. Flagged reads are almost entirely low-MAPQ (median ∼1 versus 46–52), sit on modestly shorter transcripts, and are indistinguishable in GC content. (**D**) Per-cell flag *rate* (flagged ÷ all reads) over annotated isoform count (*x*) and percent transcript covered (*y*); rate peaks at moderate isoform counts and high coverage. (**E**) Single-isoform reads split by mapping quality: bar height is the fraction of each MAPQ bin (≤3 versus *>*3) that is flagged, and the grey label gives that bin’s share of all single-isoform reads. (**F**) Flag rate by paralog status and isoform class (single, *n*_iso_ = 1; multi, *n*_iso_ ≥ 2). (**G**) Flag mechanism: interior-MAPQ residual rule (`rqr_low`) versus zero-mass deviance rule (`p0_dev_high`).

The most informative feature was transcript coverage. Reads flagged as low quality covered a *greater* fraction of their assigned reference transcript than retained or kept reads (median ∼98% versus 73–77%) despite their low mapping quality. This can be explained by reads mapping extremely well to multiple transcripts with near-complete coverage making it impossible to determine the true transcript of origin and thus get filtered out. Consistently, the per-cell flag rate peaked at transcripts with 2–3 isoforms and high coverage (up to 12.9% in HCC827 at two isoforms and 70–80% coverage (**Figure 5D**)—high coverage with modest isoform multiplicity, not single-isoform loci, carried the greatest per-read risk of flagging. On the contrary, only 1.9–3.1% of single-isoform reads had MAPQ≤3, but within that low-MAPQ minority 14.6–29.1% were flagged, versus fewer than 0.3% of the MAPQ*>*3 majority (**Figure 5E**). Single-isoform reads are therefore flagged almost exclusively when they are low-MAPQ—as expected—but such reads are rare, because unique transcripts appear to seldom produce ambiguous alignments, so few flags fall on single-isoform loci in absolute terms.

Both sources of sequence redundancy independently raised the flag rate (**Figure 5F**). First, restricting to single-isoform genes, where within-gene splicing ambiguity is absent, paralogous transcripts were still flagged roughly five-fold more often than non-paralogous ones (1.08% vs 0.21% in H1975; 2.26% vs 0.41% in HCC827; 1.29% vs 0.20% in H211), isolating between-gene paralogy as an independent driver with isoform count held fixed. Paralogous multi-isoform transcripts were flagged most of all (1.7–2.8%). Finally, flags arose through two routes—an anomalously low RQR among interior-MAPQ reads (`rqr_low`) or a MAPQ=0 read the model did not expect to be zero (`p0_dev_high`)—with the residual route dominating in every sample (66.7–77.4% of flags) (**Figure 5G**). Together these diagnostics indicate that the filter is selective rather than aggressive: it removes a small, well-defined population of ambiguously mapped reads concentrated on paralogous, multi-isoform transcripts that are highly covered by reads, while retaining the bulk of confidently mapped reads.

### 2.6 Context-aware filtering improves transcript quantification accuracy and recovery

Having established that CALMeR flags reads whose MAPQ is anomalously low given transcript and read context, we asked whether this improves downstream transcript quantification relative to fixed-threshold filtering or to probabilistic quantifiers that use no MAPQ filter at all. We compared six read-selection/quantification strategies that all begin from the *same* `minimap2` [2] transcriptome alignment and differ only in how reads are counted or assigned: (i) MAPQ ≥ 1 (the `FLAIR` [8] convention), which counts primary alignments with MAPQ ≥ 1; (ii) our proposed approach `CALMeR`, which counts the primary alignments of reads not flagged by the context-aware rule (the `quantify_counts` output of the `CALMeR` package); and four probabilistic quantifiers run over the full alignment set with no MAPQ filter — (iii) `oarfish` [10], (iv) `salmon` [9], (v) `NanoCount` [28], and (vi) `TranSigner` [29]. The first two produce integer primary-alignment counts, while `oarfish`, `salmon`, `NanoCount`, and TranSigner assign fractional abundance by expectation–maximization over the full alignment set, so all comparisons use Spearman’s *ρ*, which is invariant to the differing scales. For each strategy, we used the three PCS114 samples (H1975, HCC827, H211) to correlate the observed abundance (log_10_ scale) against the known concentration (log_10_ scale) over the Sequins Mix A panel (*n* = 160; 0.004–1,692 amol/*µ*L) and the ERCC ladder (*n* = 92), and counted the spike-in and endogenous (GENCODE) transcripts detected at ≥ 1 read.

`CALMeR` achieved the highest sequin accuracy on average and in two of the three PCS114 samples (Spearman *ρ* = 0.849, 0.842, 0.852 for H1975, HCC827, H211; **Figure 6A**), essentially tied with `NanoCount` on H1975 (both *ρ* ≈ 0.849). `NanoCount` was the next most accurate (0.849, 0.831, 0.850), followed by `oarfish` (0.829, 0.819, 0.847), `TranSigner` (0.822, 0.818, 0.846), MAPQ ≥ 1 (0.810, 0.826, 0.826), and `salmon` (0.786, 0.805, 0.838). Sequin recovery was comparable among the leading methods: `CALMeR` detected 68, 69, and 70 of 160 sequins, similar to `NanoCount` (67, 63, 72; highest in H211) and ahead of the remaining strategies (60–67). On the ERCC ladder all six strategies were indistinguishable (*ρ* ≈ 0.878, 0.868, 0.866), as expected: ERCC transcripts map uniquely at MAPQ = 60 and are unaffected by any MAPQ-based selection, providing a negative control that confirms the sequin differences are driven by the multi-mapping reads that the strategies handle differently.

**Figure 6:**
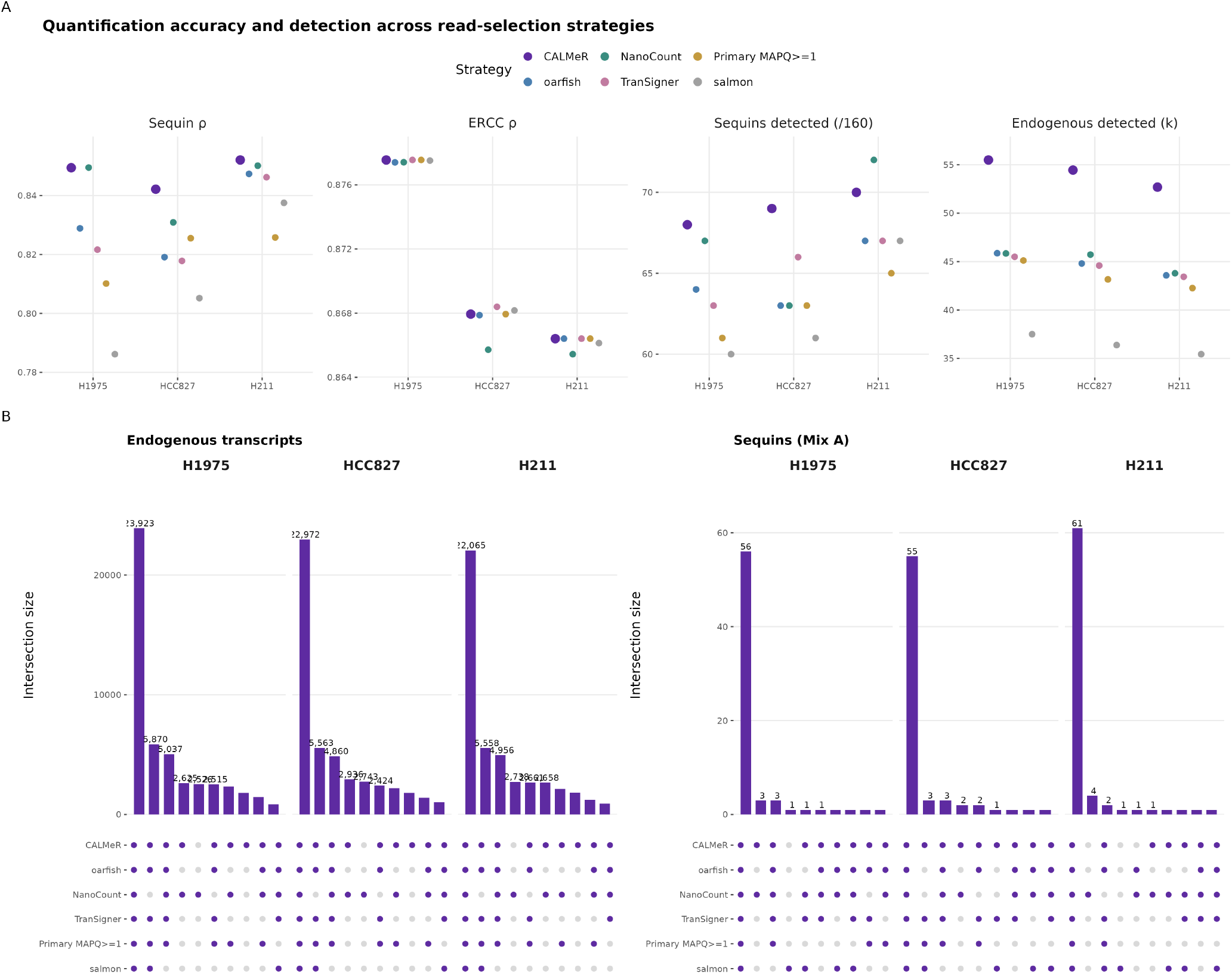
Quantification accuracy, detection, and transcript-set overlap across six read-selection/quantification strategies in three PCS114 samples (H1975, HCC827, H211). All strategies count or assign reads from the *same* `minimap2` transcriptome alignment: MAPQ ≥ 1 (primary alignments with MAPQ ≥ 1), CALMeR (unflagged primary alignments), and the probabilistic quantifiers oarfish, salmon, NanoCount, and TranSigner (expectation–maximization over all alignments, no MAPQ filter). (**A**) Accuracy and detection per strategy and sample: Spearman’s *ρ* between log_10_ observed abundance and log_10_ known concentration for Sequins Mix A and the ERCC ladder; the number of Sequins detected (of 160); and the number of endogenous GENCODE transcripts detected (thousands). (**B**) UpSet-style intersection plots of the detected-transcript sets, for endogenous transcripts (left) and Sequins Mix A (right), faceted by sample. Top bars give the size of each intersection (the ten largest per sample); the dot grid below marks which strategies participate in each intersection.

The accuracy was accompanied by a substantial gain in endogenous transcript recovery. `CALMeR` detected the most GENCODE transcripts in every sample (55,494 / 54,448 / 52,688), roughly 10,000 more than the next tier — `oarfish` (45,860 / 44,807 / 43,576), `NanoCount` (45,837 / 45,721 / 43,791), `TranSigner` (45,498 / 44,591 / 43,436), and MAPQ ≥ 1 (45,110 / 43,159 / 42,264) — and ∼18,000 more than `salmon` (37,503 / 36,379 / 35,438). The intersection structure (**Figure 6B**) indicates these extra detections are genuine rather than spurious: the large majority of transcripts are detected by all six strategies (e.g. 23,923 of the 62,510 detected in H1975), and the transcripts `CALMeR` recovers beyond the fixed-threshold and EM methods are predominantly co-detected by at least one other quantifier, leaving only a small `CALMeR`-exclusive set (∼1,800). Context-aware rescue therefore recovers a large set of transcripts that fixed-threshold and EM-based strategies discard or dilute, without loss of spike-in accuracy.

## 3 Discussion

Static MAPQ thresholds are ubiquitous in long-read RNA-seq workflows, yet the theoretical basis for any particular cutoff is weak: the threshold is chosen arbitrarily, applied uniformly across all transcript contexts, and the fraction of data discarded varies unpredictably across genes. We showed that this uniformity carries a systematic cost. MAPQ scores are not context-free signals of alignment quality, but rather they are shaped by transcript length, GC content, isoform complexity, paralog presence, and the fraction of the transcript a read covers. Applying a fixed threshold without accounting for these dependencies dis-proportionately discards reads from multi-isoform genes and longer transcripts—precisely the loci where isoform-level resolution matters most.

Rather than treating this only as a data-loss problem, we asked whether accounting for MAPQ’s context-dependence improves quantification accuracy. CALMeR reframes read selection as a per-read outlier-detection problem: it asks whether an observed MAPQ is anomalously low given the transcript and read context predicted by a fitted Zero-One-Inflated Beta regression, and retains reads whose low MAPQ is consistent with genuine biological ambiguity while removing those that are anomalous. On Sequins Mix A spike-ins, where ground-truth abundances are known, quantifying CALMeR-selected reads yielded the most accurate abundance estimates of the four strategies tested—the highest Spearman correlation with known concentration across samples —while simultaneously detecting the most transcripts. Context-aware selection therefore recovers accuracy that uniform filtering sacrifices, rather than merely trading sensitivity against specificity.

The flagged-read diagnostics clarify what CALMeR removes and why this improves rather than degrades quantification. Flagging is selective, not aggressive: only 1.2–1.6% of scorable primary reads are removed. Flagged reads are almost entirely low-MAPQ (median ≈ 1 versus 46–52 for retained reads) yet cover nearly the full length of their assigned transcript (median ≈ 98% versus 73–77%). Near-complete coverage combined with low mapping quality is the signature of reads that align almost equally well to several similar transcripts, and two independent lines of evidence confirm that redundancy—not poor alignment—drives flagging. Paralogous transcripts are flagged 1.3–1.8× more often than non-paralogous ones, and among single-isoform genes, where within-gene splicing ambiguity is absent, paralogous transcripts are still flagged roughly five-fold more often than non-paralogous ones, isolating between-gene sequence duplication as an independent driver. The removed reads are thus concentrated on exactly the loci where a single read cannot be confidently assigned to one transcript, which is where naive counting introduces the most bias.

Several limitations should be noted. The ZOIB model is fit on a random sample of 50,000 reads per sample, which reduces runtime but estimates parameters from a subset of the data. The spike-in benchmarking relies on Sequins Mix A, whose isoform complexity and transcript-length distribution are narrower than the endogenous transcriptome; the quantification biases documented here are therefore likely conservative relative to highly multi-isoform endogenous loci. CALMeR as implemented is specific to minimap2-aligned data, leveraging the near-universal adoption of minimap2 to make MAPQ interpretable across samples. Extension to other aligners would require re-characterizing the MAPQ–feature relationship for each tool.

Positioned this way, CALMeR occupies a specific niche in long-read isoform quantification: a context-aware read-selection step whose benefit is measured directly in quantification accuracy. As a standalone method it delivers high quantification accuracy—the best sequin correlation of any strategy tested—so it can be used directly to produce transcript abundances. At the same time, because its contribution is at the level of read selection rather than the assignment model itself, it is not tied to a single quantifier: the reads it retains can equally be passed to probabilistic tools that resolve isoform assignment from coverage and structural features.

## 4 Methods

### 4.1 Statistical model for context-aware MAPQ filtering

#### Response variable

For each primary alignment, the raw MAPQ score *z*_*i*_ ∈ [0, 60] was rescaled to the unit interval as 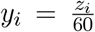 where *y*_*i*_ = 0 corresponds to fully ambiguous mapping and *y*_*i*_ = 1 to the maximum aligner-assigned uniqueness (minimap2 MAPQ = 60). Secondary and supplementary alignments were excluded from model training and scoring.

#### Distributional model

The rescaled scores *y*_*i*_ exhibit boundary inflation at both endpoints: a substantial fraction of primary alignments map with multi-isoform ambiguity (*y*_*i*_ = 0) or maximum uniqueness (*y*_*i*_ = 1), with the remainder distributed across the interior (0,1). A standard Beta regression cannot accommodate point masses at the boundaries. We therefore specified a Zero-One Inflated Beta (ZOIB) regression model, fit as the `BEINF` family in the `gamlss` framework [25]. `BEINF` parameterizes the response for alignment *i* by four quantities (*µ*_*i*_, *σ*_*i*_, *ν*_*i*_, *τ*_*i*_), each with its own linear predictor in **x**_*i*_: *µ*_*i*_ is the mean of the interior beta component, *σ*_*i*_ its dispersion, and *ν*_*i*_, *τ*_*i*_ are the (positive) mass parameters governing the boundary point masses at 0 and 1. We use the default `gamlss` links — logit for *µ* and *σ*, log for *ν* and *τ* — and hold the interior dispersion constant across alignments (*σ*_*i*_ ≡ *σ*). Writing ***β, γ, δ*** for the coefficient vectors of the *µ, ν*, and *τ* sub-models respectively,

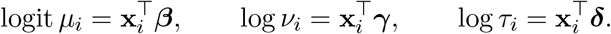

The three response regions then have the following probabilities and density, for each alignment *i* with predictor vector **x**_*i*_:

1. **Probability of full ambiguity** (*y*_*i*_ = 0, MAPQ = 0):

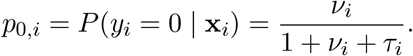
2. **Probability of maximum uniqueness** (*y*_*i*_ = 1, MAPQ = 60):

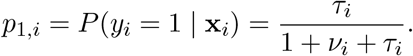
3. **Beta density for interior values** (0 *< y*_*i*_ *<* 1), carrying the remaining mass 1 − *p*_0,*i*_ − *p*_1,*i*_ = 1*/*(1 + *ν*_*i*_ + *τ*_*i*_), with mean *µ*_*i*_ and shape parameters *a*_*i*_ = *µ*_*i*_(1 − *σ*^2^)*/σ*^2^, *b*_*i*_ = (1 − *µ*_*i*_)(1 − *σ*^2^)*/σ*^2^.

To avoid degeneracy of the Beta shape parameters, *µ*_*i*_ is bounded away from {0, 1} by clipping to (10^−4^, 1 − 10^−4^). The marginal expectation under the mixture is

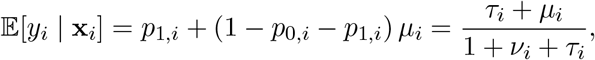

which reproduces the expected-MAPQ expression E[MAPQ | **x**_*i*_] = 60 E[*y*_*i*_ | **x**_*i*_] used for the partial-effects curves (**Figure 4**). The full coefficient vector is ***θ*** = (***β, γ, δ***); the boundary probabilities *p*_0,*i*_, *p*_1,*i*_ and the point-mass estimates 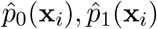 used in the outlier-scoring step below follow directly from *ν*_*i*_, *τ*_*i*_ as given above.

#### Predictor specification

Each read’s primary alignment was annotated with five covariates, drawn from the transcript annotation and the alignment itself. The covariates fall into three classes that differ in how they enter the linear predictor of the regression: smoothed (penalized B-spline on a standardized continuous variable), piecewise linear (an integer count with a kink at a fixed knot), or plain linear (a binary indicator).

- **Transcript-specific features** (constant across all reads aligned to the same transcript):
  - **Transcript length** (tx_length): total transcript length in nucleotides (sum of exon widths from the GTF). Raw values span four orders of magnitude; we applied a natural-log transform and then standardized (*z*-scored) on the training subset.
  - **GC content** (gc_content): proportion of G and C bases in the transcript sequence (FASTA), as a percent.
  - **Number of isoforms** (n_isoforms): total number of annotated isoforms for the parent gene (GTF). Counts ≥ 7 are lumped to 7 because the upper tail is sparse and very high isoform counts do not reliably distinguish ambiguity levels. The lumped count enters as a two-piece linear basis with a knot at 5: *n*_iso_ + *I*(max(*n*_iso_ − 5, 0)).
  - **Paralog presence** (has_paralog): binary indicator for whether the target transcript has at least one cross-gene paralog passing the alignment-stage filters.
- **Read-specific features** (depend on the individual alignment):
  - **Percent covered** (pct_covered): let *l*_*r*_ denote the alignment length (in nucleotides) and *l*_*t*_ the transcript length; then 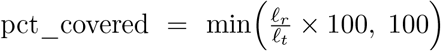 . Stated in words, this is the proportion of the transcript length spanned by the read alignment, computed as alignment_length*/*tx_length × 100 and capped at 100%. Winsorized at the 1st and 99th percentiles of the training distribution to absorb a small heavy tail at the low end, then standardized; the winsorize bounds are stored on the fitted model and reused at scoring time. Enters as a penalized B-spline.

#### Model fitting

To reduce runtime, the ZOIB model is fit on a random sample of 50,000 primary alignments drawn from each sample; the trained model is then applied to score all primary alignments in the full dataset. Since the `clean_features` function clamps rather than discards out-of-support values — lumping reads above the isoform cap into the top bin (*n*_iso_ = 7) and winsorizing percent-covered into the training range — every primary alignment carried into the model is scored.

#### Outlier scoring

For each scored alignment we compute a randomized quantile residual (RQR) [26]:

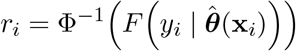

for interior values *y*_*i*_ ∈ (0, 1), with the analogous randomization over the boundary CDF interval for *y*_*i*_ ∈ {0, 1}. Under correct model specification, *r*_*i*_ is approximately a standard normal. Deviations from *N* (0, 1) jointly summarize model adequacy and per-read anomaly. We additionally record the predicted point-mass probabilities

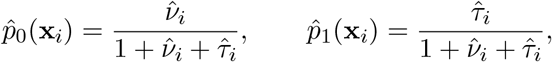

which drive the point-mass component of the flagging rule below.

#### Flagging criterion

Although the RQR is well defined for every *y*_*i*_, the residual is randomized over a CDF interval at the ZOIB point masses (*y*_*i*_ = 0 corresponding to MAPQ = 0 and *y*_*i*_ = 1 to MAPQ = 60). Therefore, a single draw at *y*_*i*_ ∈ {0, 1} cannot reliably distinguish model-expected from anomalous point-mass outcomes [26]. Following [30], who recommend handling point-mass observations separately rather than folding them into a unified RQR-tail rule in zero-adjusted regression models, we apply an asymmetric flagging criterion that pairs a residual-based test on the interior part with a deviance-based test on the lower mass:

- **Interior part** (*y*_*i*_ ∈ (0, 1), i.e. MAPQ ∈ {1, …, 59}): flag if the RQR falls below a robust lower threshold,

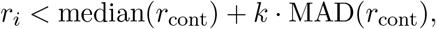

where the median and MAD are computed on the interior-MAPQ reads only. We use *k* = −2 by default.
- **Lower point mass** (*y*_*i*_ = 0, MAPQ = 0): flag if the Bernoulli deviance lies in the upper tail of the zero-mass deviance distribution,

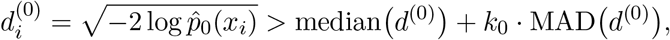

where the median and MAD are computed over the MAPQ= 0 reads and *k*_0_ = 2 by default. Like the interior rule, this is a robust (median/MAD) threshold estimated per sample from the empirical deviance distribution rather than a fixed probability cutoff. A large deviance means the model assigned the read little probability of landing at the zero mass; this catches reads with strong covariate support (high coverage, low isoform count, no paralogous context) whose alignment nevertheless collapsed to MAPQ = 0.
- **Upper point mass** (*y*_*i*_ = 1, MAPQ = 60): never flagged. MAPQ= 60 is the maximum achievable mapping quality and cannot be “unusually low” under any context.

Flagged reads, together with all secondary and supplementary alignments sharing their read ID, are removed from the output BAM.

### 4.2 Paralog Detection

For each annotated transcript, a binary indicator `has_paralog` ∈ {0, 1} records whether the transcript has at least one cross-gene paralog passing a set of length, coverage, and identity filters. This computation runs once per transcriptome annotation and is independent of any individual sample’s reads. The transcriptome FASTA was aligned against itself using `minimap2` [2] with parameters `-D -x asm20 –secondary=yes –N 100 -c`. The `-x asm20` preset is tuned for sequence pairs with up to roughly 20% divergence and imposes an effective ∼ 80% identity floor at the alignment stage. The `-D` flag suppresses trivial self-mappings (a transcript aligning to itself by name) without affecting the dual direction of any other pair. We deliberately did *not* use the alternative `-X` flag (equivalent to `-D –dual=no`): `-X` also suppresses one direction of every transcript pair, so that for a paralog pair (*A, B*) `minimap2` reports the alignment in only one direction. Combined with our per-query aggregation step (below), `-X` silently fails to flag whichever member of each pair was emitted only as the target.

For each resulting alignment record in the resulting Pairwise mApping Format (PAF) file, we computed:

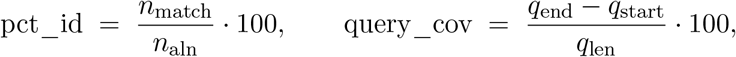

where *n*_match_ (PAF column 10) is the number of matching bases and *n*_aln_ (column 11) is the alignment block length (including gaps and mismatches) reported by `minimap2`; *q*_len_ (column 2) is the total length of the query transcript, and (*q*_start_, *q*_end_) (columns 3 and 4) are the 0-based half-open start and end coordinates of the alignment along the query.

#### Filtering

An aligned read was retained as a qualifying paralog hit only if:

1. the query and target transcripts had distinct identifiers (a defensive check; `-D` already removes most such rows);
2. *n*_aln_ ≥ 50 base pairs;
3. query_cov ≥ 70%; and
4. the query and target transcripts mapped to different `gene_id` values in the GTF annotation, where this mapping was obtained from the same GTF used elsewhere in the pipeline.

The minimum alignment length drops short chain artefacts. The query-coverage threshold remains important even given the `asm20` identity floor: `asm20` constrains only the identity of the bases that did align, not what fraction of the query participated. Without the 70% threshold, two genes that happen to share a single high-identity conserved domain (for example, a ∼ 80 bp DNA-binding motif within otherwise unrelated 1500 bp transcripts) would survive identity filtering despite being unrelated outside the domain. Requiring ≥ 70% query coverage enforces whole-transcript homology, which is the relevant notion of paralog in transcript terms. The final cross-gene criterion excludes alternative isoforms of the same gene, which would otherwise dominate the set of high-identity hits.

#### Per-transcript aggregation

For each annotated query transcript *t*, let *H*_*t*_ denote the set of cross-gene alignments passing the filters above. We defined the binary paralog indicator as

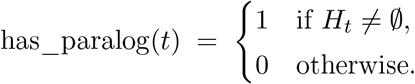

Transcripts with no qualifying cross-gene hit therefore receive has_paralog = 0.

We chose a binary indicator rather than a continuous identity score (max_*h*∈*H*_*t* pct_id(*h*)) because the `asm20` preset’s ∼ 80% identity floor renders the post-filter pct_id distribution *bimodal* : a point mass at zero (transcripts with no qualifying hit) and a cluster at 80–100% (transcripts with at least one), with essentially no observations in between. A linear coefficient on such a bimodal variable would conflate the between-cluster shift with a within-cluster slope that the data cannot identify; the binary indicator captures the same information without that confound, and a penalized smoother is not meaningful on a two-valued covariate. The alignment-stage filters (`asm20` + the length, coverage, and cross-gene requirements) together encode the operative paralog cutoff.

#### Integration into the read-level model

At feature-extraction time we joined the per-transcript paralog table onto every primary alignment by `transcript_id` (with a version-stripped fallback to accommodate annotation-version mismatches), so that each read inherits the binary paralog flag of its assigned transcript. Reads on transcripts not in the paralog table inherit has_paralog = 0. The flag enters the ZOIB model as a single linear coefficient in each of the three sub-models (interior, zero-inflation, one-inflation), with no smoother and no standardization.

### 4.3 Long-read RNA-seq benchmark datasets

#### Dataset selection and scope

To characterize the relationship between MAPQ and transcript-level features across diverse long-read RNA-seq conditions, and to benchmark the impact of MAPQ filtering strategies on quantification, we identified five publicly available datasets from recent large-scale bench-marking studies. Together these span direct RNA, direct cDNA, PCR cDNA, R2C2, and CapTrap-Seq library preparations across eight sequencing kits (**Table S1**): (i) Dong et al. [17], (ii) the Long-read RNA-Seq Genome Annotation Assessment Project (LRGASP) [14], (iii) LongBench [16], (iv) the Singapore Nanopore Expression (SG-NEx) Project [15], and (v) [18], who evaluate CapTrap-Seq against nanopore direct RNA and other non-nanopore protocols. All five datasets include synthetic spike-in controls — Lexogen SIRVs [19], Sequins [20], or a combination of both — with composition and concentration varying by sample (**Table S2**).

#### Unified preprocessing pipeline

To enable cross-benchmark comparisons, all samples were processed through a single standardized pipeline. Raw FASTQ files were quality-filtered to retain only reads with mean base quality *Q* ≥ 10. All samples were aligned to a common reference: GENCODE v46 (GRCh38) for endogenous transcripts, with spike-in annotations specific to each benchmark downloaded and merged into the GENCODE GTF to produce comprehensive per-benchmark annotation files. Transcriptome sequences were extracted from the merged GTF using `gffread` to generate FASTA files for transcriptome-based alignment [31]. Quality-filtered reads were aligned to the transcriptome using `minimap2` [2]. Gene-isoform level quantification was performed using `oarfish` [10].

#### Spike-in benchmarking subset

To obtain ground-truth quantification measurements for evaluating the impact of MAPQ filtering on accuracy, we focused on eight PCR cDNA samples carrying Sequins Mix A v1 [21] from Dong et al. [17] and LongBench [16] (three PCS109 samples and five PCS114 samples, respectively). These samples were selected because they span the broadest dynamic range of transcript-level features among the available spike-in panels: isoform count per gene 1–4, transcript length 283–6,943 bp, and GC content 32.5–68.9%. For a given MAPQ filter threshold *α*, quantification bias was assessed as the change in the log_2_-ratio of observed to expected spike-in proportions relative to the no-filter baseline, Δ log_2_ FC_*t*_(*α*), defined in Equations 1–2. In an unbiased setting, Δ log_2_ FC = 0; systematic deviations indicate filter-induced representation bias.

### 4.4 Comparison to existing methods

#### Alignment and quantification settings

All strategies started from a single transcriptome alignment produced with `minimap2 --eqx -N 100 -ax map-ont` against the merged GENCODE v46 + spike-in transcriptome FASTA. oarfish was run as `oarfish -a <bam> --filter-group no-filters` `--model-coverage` and salmon as `salmon quant -t <ref> --libType A -a <bam>`, both consuming the same no-filter BAM. NanoCount (v1.1.0) was run as `NanoCount -i <bam> -o <out.tsv>` `--extra_tx_info`, taking its estimated read counts (`est_count`) as abundances. TranSigner (v1.1.3) was run in two stages on a read-name–sorted copy of the same BAM: first `transigner pre -i` `<name_sorted.bam> -d <out>` to build the read–transcript compatibility scores, then `transigner em` `-s scores.csv -d <out> -u unmapped.txt -m tmap.csv -dtype ont_cdna -n 100` to estimate per-transcript abundances. CALMeR and MAPQ ≥ 1 counted primary alignments from that BAM.

### 4.5 Code and software availability

Code to reproduce all preprocessing, analyses, and figures in this manuscript is available on GitHub at https://github.com/sparthib/calmerpaper. CALMeR is available as an open-source R package on GitHub at https://github.com/sparthib/CALMeR

## 4.6 Acknowledgments, Funding, Authorship Contributions

### Use of AI tools

During the preparation of this manuscript, the authors used a large language model (LLM) Claude Opus 4.8 and GPT-6.1 Sol, to assist with generating and debugging analysis code and to edit the language of the manuscript for clarity and concision. The authors reviewed and verified all code output and edited text, and take full responsibility for the content of the publication.

## Acknowledgments

We acknowledge the Joint High Performance Computing Exchange (JHPCE) at the Johns Hopkins Bloomberg School of Public Health for computational support, and members of the Hicks lab for their valuable feedback.

## Author Contributions Statement

**SP:** Software, Data curation, Methodology, Formal analysis, Investigation, Visualization, Writing – original draft, Writing – review & editing; **SCH:** Supervision, Funding acquisition, Project administration, Software, Writing – original draft, Writing – review & editing.

## Supplementary Materials

### Supplementary Figures

**Supplementary Figure S1:**
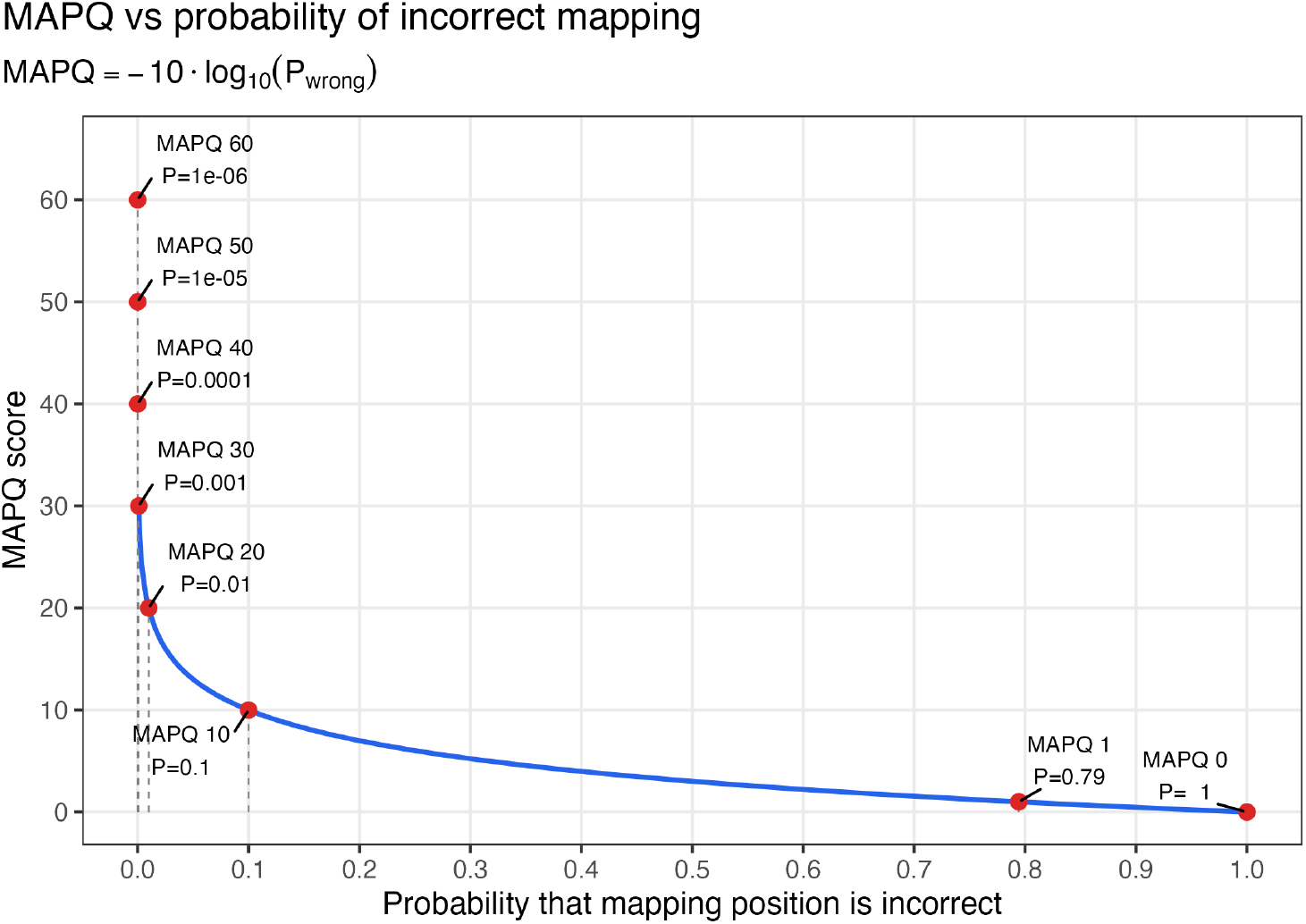
Relationship between MAPQ score (*y*-axis) and the probability that a mapping position is incorrect, Pr{wrong} = 10^−MAPQ*/*10^ (*x*-axis). Red points mark MAPQ values 0, 1, 10, 20, 30, 40, 50, and 60, which are common thresholds to remove reads below a given value, such as a MAPQ of 30.

**Supplementary Figure S2:**
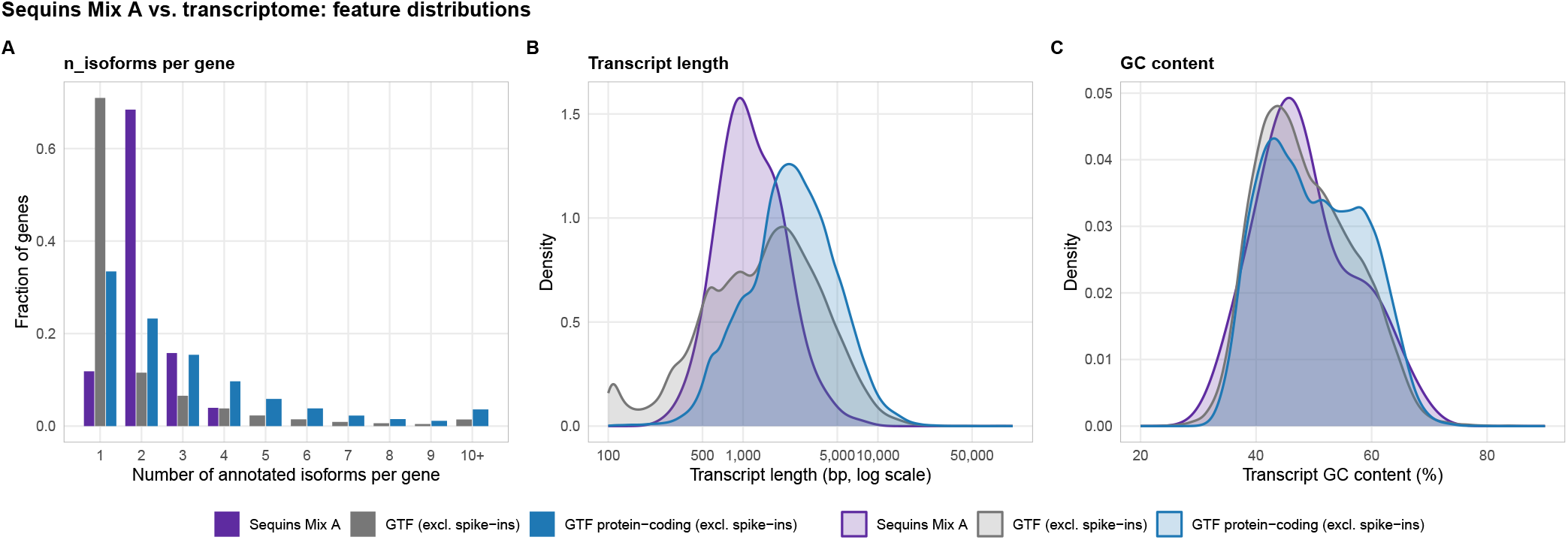
Sequins Mix A feature distributions versus the transcriptome. **(A)** Over-lapping per-category densities for number of annotated isoforms per gene, **(B)** transcript length (log scale), and **(C)** transcript GC content.

**Supplementary Figure S3:**
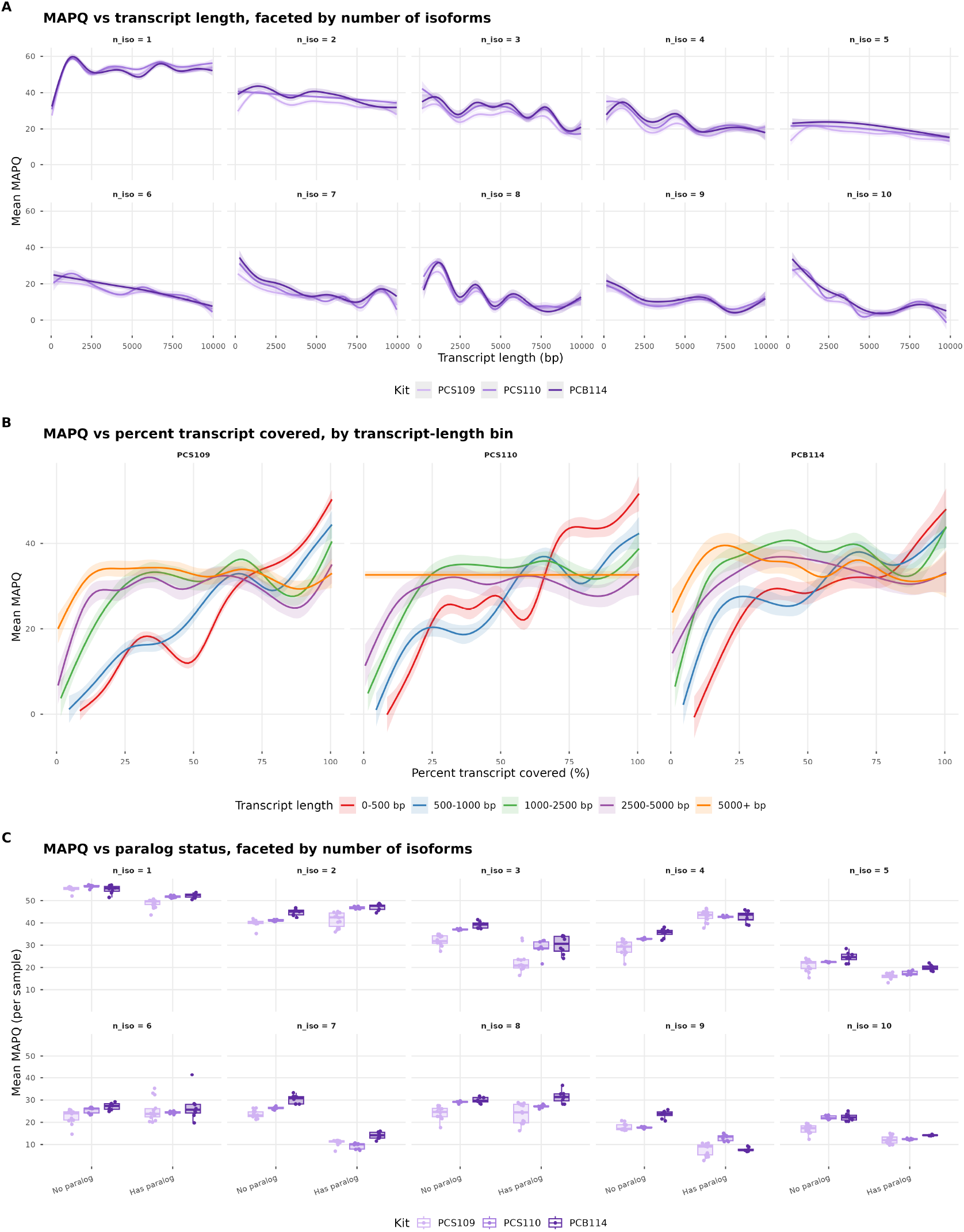
The complex landscape of MAPQ between the interaction of isoform count, transcript coverage, transcript coverage, and paralog status. All panels use primary alignments from PCR cDNA kits PCS109, PCS110, and PCS114 and calculate per sample mean MAPQ for a given strata. (**A**) Mean MAPQ versus transcript length in 500 bp bins (up to 10,000 bp), stratified by annotated isoform count (1–10). Lines and ribbons show kit-specific means ±1 standard error across samples. (**B**) Mean MAPQ versus percent transcript covered (or fraction of reference transcript length), calculated at 1% resolution and averaged across samples, by kit and transcript-length class (0–500, 500–1000, 1000–2500, 2500–5000, 5000+ bp). Curves show smoothed Generalized Additive Models (GAM) with 95% confidence intervals. (**C**) Mean MAPQ for transcripts with or without cross-gene paralogs, stratified by isoform count. Boxplots summarize per-sample distributions (outliers suppressed); points are coloured by kit.

**Supplementary Figure S4:**
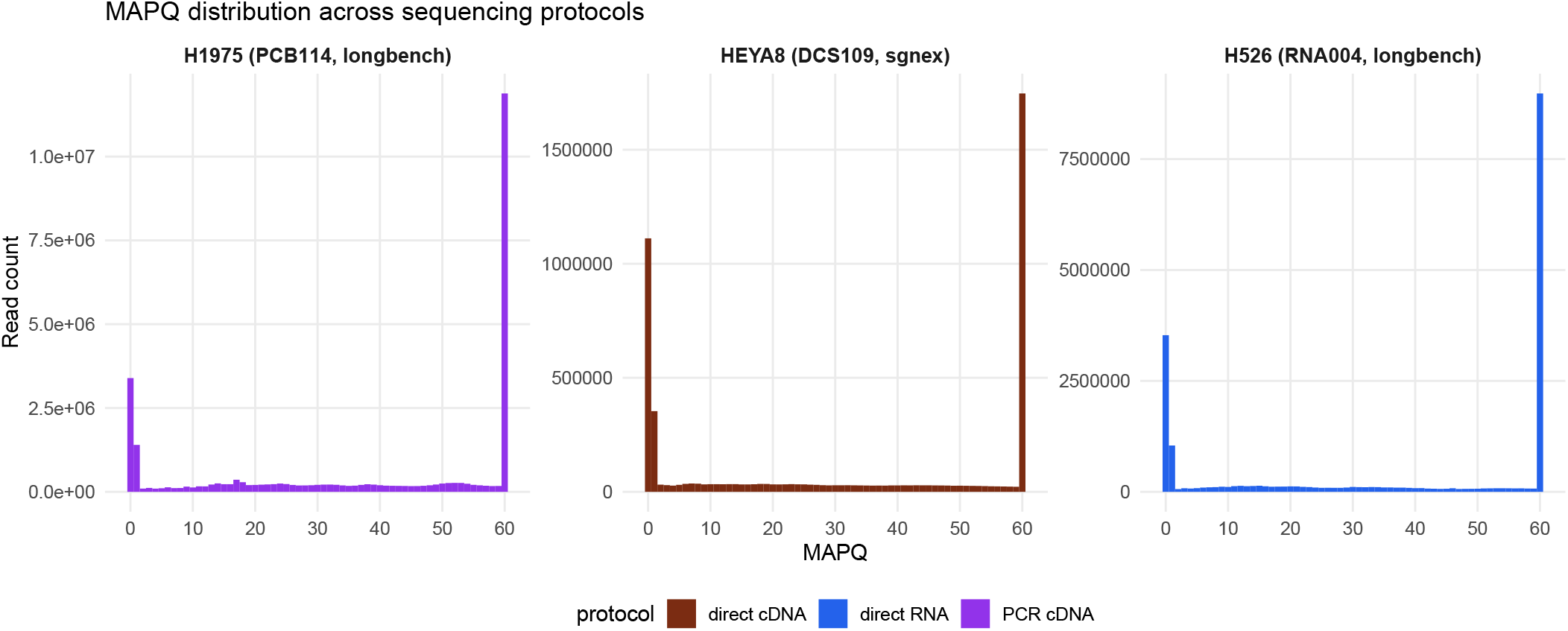
MAPQ distribution varies with sequencing protocol. Per-read MAPQ histograms for one representative sample from each of three long-read sequencing protocols: **(A)** PCR cDNA, **(B)** direct RNA, and **(C)** direct cDNA. Each panel title lists the cell line, library-preparation kit, and source benchmark dataset for the displayed sample. Bars are coloured by protocol; the *y*-axis is read count and is free across panels because absolute read depth differs between samples. The characteristic bimodal shape (mass at MAPQ = 0 and MAPQ = 60) is present across all three chemistries.

**Supplementary Figure S5:**
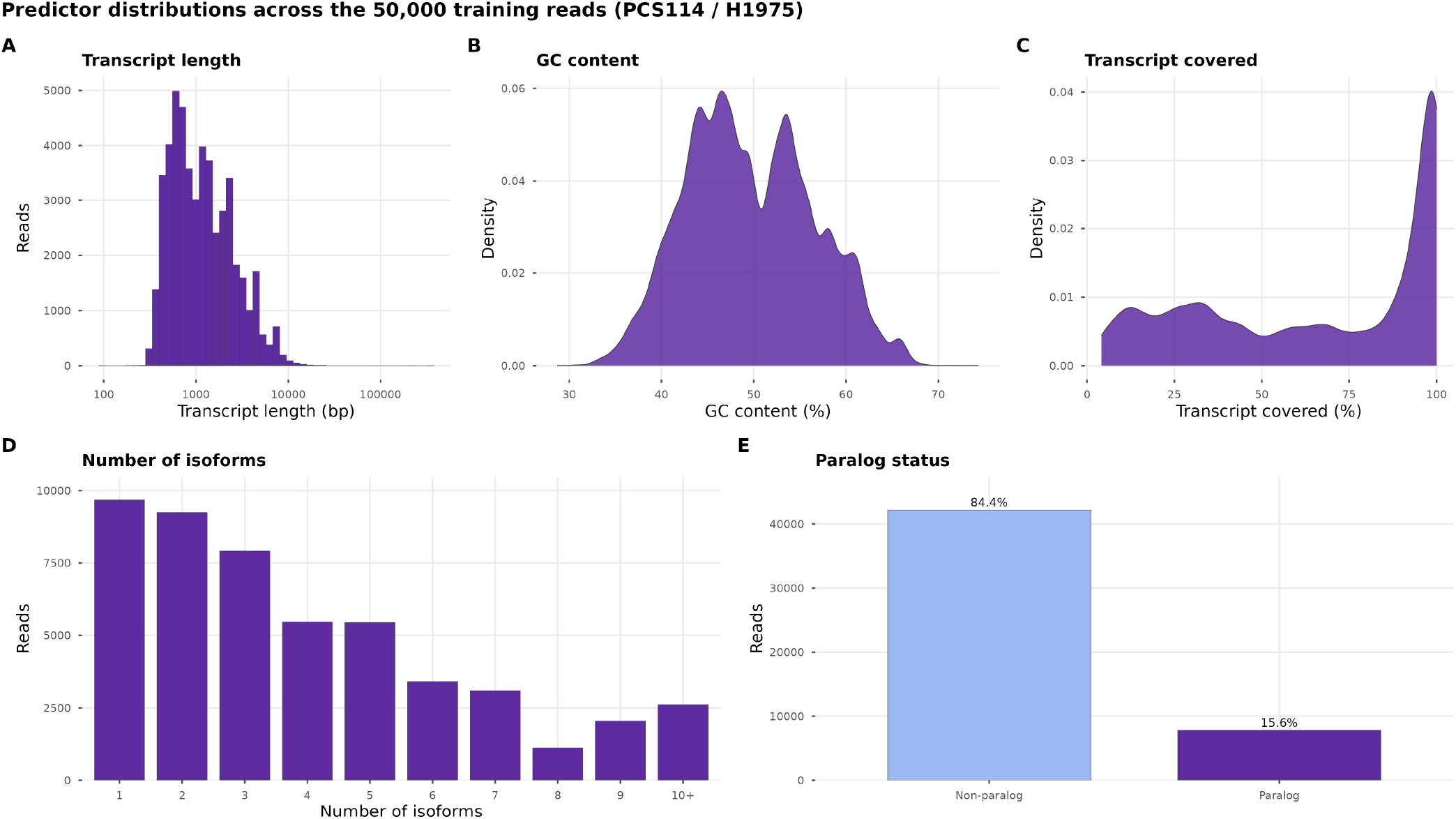
Distributions of the five model predictors on the random subset of *n* = 50,000 primary alignments used to fit the BEINF model for the PCS114 / H1975 sample (GSM9135480). Values reflect the canonical preprocessing applied by `clean_features()`: **(A)** `n_isoforms` is lumped at 7 (counts ≥ 7 are clamped to 7); **(B)** `tx_length` is shown on a log-bp x-axis after the natural-log transform; **(C)** `gc_content` is plotted unchanged; **(D)** `pct_covered` is winsorized at the 1st and 99th percentiles of the training sample; **(E)** `has_paralog` is a binary indicator for at least one cross-gene paralog passing the alignment-stage filters. Histograms are used for the discrete or count-like predictors (`n_isoforms`, `tx_length`, `has_paralog`) and kernel density estimates for the continuous percentage variables (`gc_content`, `pct_covered`).

**Supplementary Figure S6:**
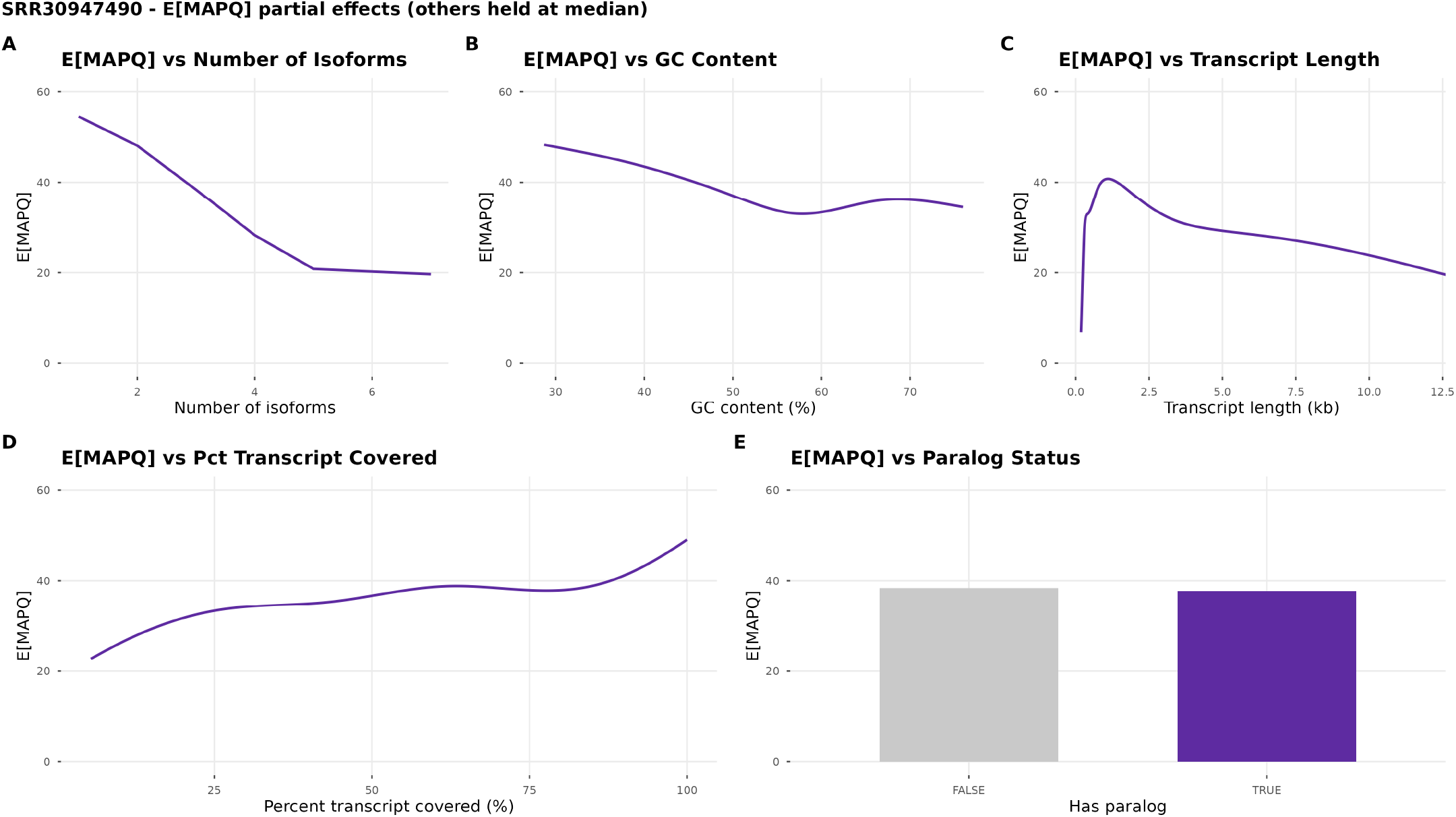
Partial effects of each predictor on expected MAPQ, direct-RNA (RNA004). Direct-RNA counterpart of Figure 4, with the BEINF model refit on the RNA004 H1975 sample (Sequins Mix A; the direct-RNA analogue of the PCR H1975 sample). Each panel varies one predictor across its training range while holding the other predictors at their median value (or FALSE for the binary `has_paralog` indicator), and plots E[MAPQ] = 60 (*τ* + *µ*)*/*(1 + *ν* + *τ* ) under the resulting prediction. (A) `n_isoforms` is entered as a two-piece linear basis with a knot at 5. (B–D) The smoothed predictors `gc_content`, `tx_length` (log-bp *x*-axis), and `pct_covered` are entered through penalized B-splines and appear as continuous curves. (E) `has_paralog` is binary and shown as a column comparison. The partial-effect trends mirror those of the PCR cDNA model (Figure 4), indicating the MAPQ–feature relationships are not specific to the PCR preparation.

**Supplementary Figure S7:**
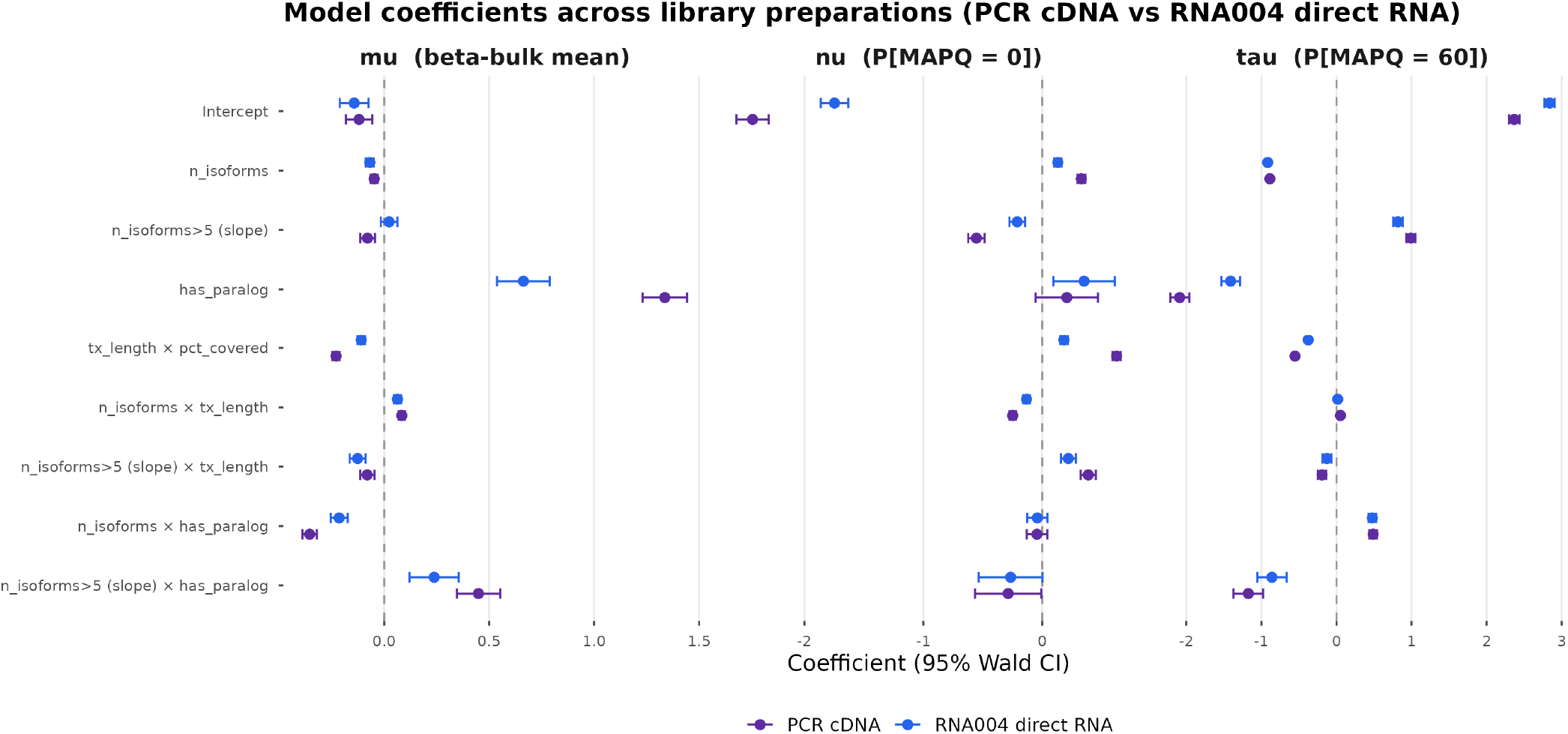
Model coefficients are consistent across library preparations (PCR cDNA vs. RNA004 direct RNA). Parametric coefficients (points) with 95% Wald confidence intervals (horizontal bars) for the BEINF MAPQ model, overlaid for the two preparations of the H1975 cell line: PCR cDNA (`GSM9135480`, Sequins Mix A, purple) and RNA004 direct RNA (`SRR30947490`, blue). Both models use the same forward-stepwise, BIC-selected specification (penalized B-splines for the three continuous covariates, all five main effects, and all three interactions; *µ, ν*, and *τ* modeled, with *σ* ∼ 1), refit independently on a 50,000 complete-case training subset of each sample. Panels correspond to the three modeled distribution parameters: *µ* (beta-bulk mean of MAPQ*/*60), *ν* (P[MAPQ = 0]), and *τ* (P[MAPQ = 60]). Penalized-spline basis terms are not interpretable point coefficients and are omitted; the displayed terms are the two-piece `n_isoforms` basis (with a knot at 5), `has_paralog`, the three interaction terms, and the intercept. The dashed vertical line marks a null coefficient of zero. The largely overlapping intervals between the two preparations indicate that the MAPQ–feature relationships learned by the model are not specific to the PCR protocol.

### Supplementary Tables

**Supplementary Table S1:** Summary of spike-in datasets. Number of FASTQ files (runs) per protocol/kit combination across the five public benchmark datasets.

| # | **Protocol** | **Kit** | **CapTrap-seq** | **Dong et al.** | **LongBench** | **LRGASP** | **SG-NEx** | **Total** |
|---|---|---|---|---|---|---|---|---|
| 1 | captrap | captrap_linker | 2 | 0 | 0 | 6 | 0 | 8 |
| 2 | PCR cDNA | PCS109 | 0 | 6 | 0 | 0 | 8 | 14 |
| 3 | PCR cDNA | PCS110 | 0 | 0 | 0 | 9 | 0 | 9 |
| 4 | PCR cDNA | PCS114 | 0 | 0 | 8 | 0 | 0 | 8 |
| 5 | R2C2 | R2C2 | 0 | 0 | 0 | 18 | 0 | 18 |
| 6 | direct RNA | RNA002 | 2 | 0 | 0 | 12 | 14 | 28 |
| 7 | direct RNA | RNA004 | 0 | 0 | 8 | 0 | 0 | 8 |
| 8 | direct cDNA | DCS109 | 0 | 0 | 0 | 0 | 12 | 12 |
|  |  | **Total** | 4 | 6 | 16 | 45 | 34 | 105 |

**Supplementary Table S2:** Spike-in datasets by benchmark, spike-in mix, protocol and kit. Counts are number of FASTQ files (runs).

| **Benchmark** | **Spike-in mix** | **Protocol** | **Kit** | **n** |
|---|---|---|---|---|
| CapTrap-seq | ERCC, SIRV1 E1, SIRV1 E2 | captrap | captrap_linker | 2 |
| CapTrap-seq | ERCC, SIRV1 E1, SIRV1 E2 | direct RNA | RNA002 | 2 |
| Dong et al. | Dong sequin Mix A | PCR cDNA | PCS109 | 3 |
| Dong et al. | Dong sequin Mix B | PCR cDNA | PCS109 | 3 |
| LongBench | Dong sequin Mix A , SIRV-4 | PCR cDNA | PCS114 | 5 |
| LongBench | Dong sequin Mix B , SIRV-4 | PCR cDNA | PCS114 | 3 |
| LongBench | Dong sequin Mix A , SIRV-4 | direct RNA | RNA004 | 5 |
| LongBench | Dong sequin Mix B , SIRV-4 | direct RNA | RNA004 | 3 |
| LRGASP | SIRV-4 | captrap | captrap_linker | 6 |
| LRGASP | SIRV-4 | PCR cDNA | PCS110 | 9 |
| LRGASP | SIRV-4 | R2C2 | R2C2 | 18 |
| LRGASP | SIRV-4 | direct RNA | RNA002 | 12 |
| SG-NEx | SGNex sequin Mix A V1 | PCR cDNA | PCS109 | 2 |
| SG-NEx | SIRV-4 | PCR cDNA | PCS109 | 3 |
| SG-NEx | SGNex sequin Mix A V2, SIRV-1 E2 | PCR cDNA | PCS109 | 3 |
| SG-NEx | SGNex sequin Mix A V1 | direct cDNA | DCS109 | 2 |
| SG-NEx | SGNex sequin Mix A V1 | direct cDNA | DCS109 | 4 |
| SG-NEx | SIRV-4 | direct cDNA | DCS109 | 3 |
| SG-NEx | SGNex sequin Mix A V2, SIRV-1 E2 | direct cDNA | DCS109 | 3 |
| SG-NEx | SGNex sequin Mix A V1 | direct RNA | RNA002 | 4 |
| SG-NEx | SIRV-4 | direct RNA | RNA002 | 5 |
| SG-NEx | SGNex sequin Mix A V2 , SIRV-1 E2 | direct RNA | RNA002 | 5 |
|  |  |  | **Total** | **105** |

